# Breedbase 2.0 - expanding the functionality for traditional breeding and transgenics

**DOI:** 10.64898/2026.09.28.755108

**Authors:** Srikanth Kumar Karaikal, Ryan Preble, David J Waring, Christiano C Simoes, Isaak Y Tecle, Naama Menda, Titima Tantikanjana, Jing Zhang, Katherine Eaton, Keo Corak, Clayton Birkett, Allison Krill Brown, Mitchell Feldmann, Christian Lamm, Mercedes Thieme, Prasad Peteti, Heather Manching, Adrian Powell, Jean-Luc Jannink, Lukas A Mueller

## Abstract

Breedbase (https://breedbase.org/) is a comprehensive, web-based, open source breeding management and analysis software for plant breeding that was introduced to address gaps with existing software in dealing with modern plant breeding data. Breedbase is continually improved and developed, with new features being added as required by the breeding community. Since its inception, the capabilities of Breedbase were improved to better serve the diverse data modeling needs of additional crops beyond our original ensemble of cassava, banana, yam, sweet potato, tomato, rice, and wheat. We describe here major updates that have been made in the past 4 years, including: (1) improved handling of repetitive measurements; (2) a redesigned experimental treatments interface; (3) a more fine-grained user permission system; (4) improved trial quality control workflows; (5) intercropping support; (6) the addition of a “decision meeting” feature that helps decision makers make the best selection with the available data; (7) transgenics and tracking tools and finally, (8) an improved field layout tool which consolidates several trials in one field. These new features integrate seamlessly with the previous feature set of Breedbase - making it one of the most feature rich breeding management softwares available.

## Introduction

Modern plant breeding relies increasingly on a combination of genomic and high throughput phenotyping technologies of a large number of plant accessions, and thus has become an immensely data intensive undertaking. By contrast, a traditional phenotypic breeding approach would involve only using parental accessions to produce a population of intercrosses from which promising progeny were selected for further crosses, based solely on phenotypic data (Breseghello and Coelho 2013). Where previous breeding methodologies involved statistical modeling of phenotypic data on the field scale (Wallace et al. 2018), modern workflows and newer breeding paradigms such as genomic selection and prediction require vast volumes of data pulled from many locations and seasons, precise field randomizations, systematic phenotype collection, and consistent genome sequencing data (Crossa et al. 2017). Modern genomic selection (GS) adds an additional layer of complexity: rather than using direct phenotypic data to make selection decisions, breeding values derived from genotype-based predictions are used to select accessions for further breeding (Ribaut and Hoisington 1998; Jannink et al. 2010).

GS breeding has advantages over traditional breeding due to the reduced dependence on high and low throughput phenotyping, which is often labor intensive and expensive compared to efficient and inexpensive modern genotyping methods. Once a sufficient volume of data has been collected initially, a model based on genotypic and phenotypic data can be used to calculate breeding values, or a predictive score over the worth of a given germplasm in the breeding lifecycle (Meuwissen and Goddard 2001). However, this upfront cost in data volume and complexity can be prohibitive in small breeding programs, and are hard to manage using local data storage in spreadsheets. This issue is especially relevant in developing countries, which stand to benefit the most from accelerated plant breeding to buttress food production while also having the largest economic barriers to adopting modern genomic breeding. Breedbase (Morales et al. 2022) was introduced as a tool and digital ecosystem for managing these complex data, solving a key infrastructure problem in plant breeding.

Breedbase is a web-based breeding management and analysis system that has been designed to organize the management, collection, and analysis of breeding data in a digital ecosystem in which all data are only entered and processed digitally, with the selection process and required analyses tightly integrated in the system. Breedbase provides a single platform to upload genetic and phenotypic data to build modern genomic breeding models. Breedbase is integrated with Field Book (https://github.com/PhenoApps/Field-Book) (Rife and Poland 2014), an open source Android app for the collection of agricultural phenotypic data, and Breedbase provides native support for uploading genotyping protocols, transformation vectors, high-throughput phenotyping data like near infrared spectroscopy (NIRS), and metabolomics (Hershberger et al. 2021). By utilizing Field Book to capture data and Breedbase to store and analyze the data, users have the ability to fully digitize their breeding workflow with intuitive app interfaces and the power of state of the art data analytics. This enhances data quality and significantly reduces technical complexity for end users. The usage of relational database techniques facilitates data management, such as minimizing data duplication, enhancing the productivity of plant breeding programs (Cobb et al. 2019). Through a standardized Breeding Application Programming Interface (BrAPI) (Selby et al. 2019; Selby et al. 2025), Breedbase can interact with many other tools and analysis software such as Mr. Bean (Aparicio et al. 2024), expanding the capabilities of the digital ecosystem almost infinitely. Breedbase is developed under an open source model, with the source code being available through a collection of repositories hosted on GitHub (https://github.com/solgenomics). For ease of deployment, a Docker container is also made available with a completely installed, configured and tested system that can be downloaded from the Docker Hub site (https://hub.docker.com/u/breedbase/). Breedbase is accompanied by a user manual detailing the usage of the site, configuration options, and how to host a new instance.

Since the publication of Breedbase 1.0 in 2022, Breedbase has expanded its capabilities to make it more useful and accessible to a growing ensemble of breeding programs and accurately model a wider array of breeding experiments. The needs of a strawberry breeding program (Feldmann et al. 2024) necessitated the introduction of more advanced phenotypic data modeling that supported collecting repetitive trait measurements. To aid in both the rigorous tracking of experiment data and breeding decision making, we incorporated a system of tracking decisions to advance or retire accessions through the breeding lifecycle. Trials with intercropped accessions and more complicated experimental designs prompted the addition of more flexible treatment applications and storing multiple accessions in relation to a plot. Accounting for the effects of location on trial data necessitated visualizing phenotypic data in trials that are replicated in different places. Transgenic studies required the addition of tools to model plant transformations. These improvements and the future directions for Breedbase are discussed here.

## Experimental Treatments

Breedbase 1.0 featured the ability to design trials with experimental treatments. When different growth scenarios are evaluated, it is important to specify the exact conditions every plot, subplot, plant, or tissue sample experiences. This is useful for evaluating effects of drought and watering regimes, fertilizers, crop protection, and many other use cases. These conditions feed forward to statistical analyses and breeding decisions, and so must be accurately modeled in the database. However, Breedbase 1.0 had significant limitations. Only plots could receive a treatment in most cases, and subplots could only receive individual treatments, not treatment combinations. Furthermore, users could only design treatments with binary values – plots could only be designated as having received or having not received a treatment. For experimental treatments with multiple application levels, repeated applications, or other treatments that could not be accurately described as a simple binary, users were forced to employ one-hot encoding to store the treatment values. For example, a user that designed a trial with three levels of fertilizer – low, medium, and high – would have to create three separate treatments, called *fertilizer_low, fertilizer_medium*, and *fertilizer_high*. In the trial design spreadsheet, the user would then enter a 0 if that treatment level was not present in the plot of that row, or 1 if the treatment level was present. The same mechanism had to be used if the same treatment was applied multiple times over the course of the study, with each treatment column referring to the same treatment at a different application time.

In the database backend, these treatments were stored as virtual projects named with the parent trial name followed by the name of the treatment column. These virtual treatment projects contained a list of plots to which their treatment was applied. Users had the option of accessing and adding metadata to these trials, such as detailed descriptions of the treatments being applied. While this was sufficient to model the application of experimental treatments to a field trial, this approach meant individual treatment applications were never stored with a robust, reproducible, relational-database approach. Two identical treatment names across two different trials had no intrinsic connection to one another. Should the same treatment be applied identically in different experiments, there was no database record confirming that they were indeed the same treatment, and storage space was wasted saving the same treatment multiple times. Additionally, searching for trials based on the treatments applied in those trials was neither programmatic nor pragmatic.

Breedbase 1.0 also featured the ability to describe the management regimes of trials. Individual management factors describe how a field trial was managed as a whole, such as irrigation regimes, pesticide applications, tilling practices, or other management practices whose sum create the trial’s management regime. Somewhat confusingly, however, Breedbase 1.0 made no distinction between management factors and experimental treatments. Management factors, similarly to experimental treatments, were stored as virtual projects with lists of plots to which a management factor was applied. The only difference between the two was semantic: experimental treatments were applied to a subset of the plots in a trial while management factors were applied to all plots in a trial. Indeed, Breedbase 1.0 didn’t even distinguish treatments from management factors in the user interface: the terms were used interchangeably, and adding management factors to a trial’s management regime was functionally the same as adding experimental treatments to a trial.

To address these issues, Breedbase 2.0 overhauled the handling of treatments and management regimes. Management regimes have been redesigned to include a type (configurable per instance as in Breedbase 1.0), a description, an application schedule, start and end dates, and records of completion. These data are now stored as JSON strings in association with individual trials. The user experience has been streamlined so that the management regime is accessible from the trial detail page, no longer necessitating traveling to a new page to view and edit field management data.

Experimental treatments are now modeled with the same structure as phenotypic data. Each treatment application has an associated observation unit (plot, subplot, plant, tissue sample, etc.), a timestamp, and a possible value determined by the individual treatment whose behavior is recorded and standardized with an experimental treatment ontology. As with trait ontologies, treatment ontologies define a hierarchy of allowable treatments that can be applied to stocks along with their data type, acceptable ranges, and whether these treatments can be applied repetitively. Configurable on a per-instance basis, users may also choose to design their own treatment ontologies for use with their crop system. This enables the experimental structure of the field trial, much like phenotypic collections, to be rigorously controlled and comparable across many trials. One-hot encoding is no longer necessary, and users can upload treatment data in exactly the same way as phenotypic data, streamlining the process of storing trial data on Breedbase. With Breedbase 2.0, treatment values can be displayed on custom printed labels directly from the Label Designer for quick identification of what treatments were applied to a given stock. Furthermore, expanding treatments to be applicable to all stocks, not just plots, enables much finer control over the experimental design. Treatments can be applied to subplots, individual plants, or even individual tissue samples, allowing agronomists to model sophisticated experiments that were not well supported in Breedbase 1.0 (Figure 1).

**Figure 1:**
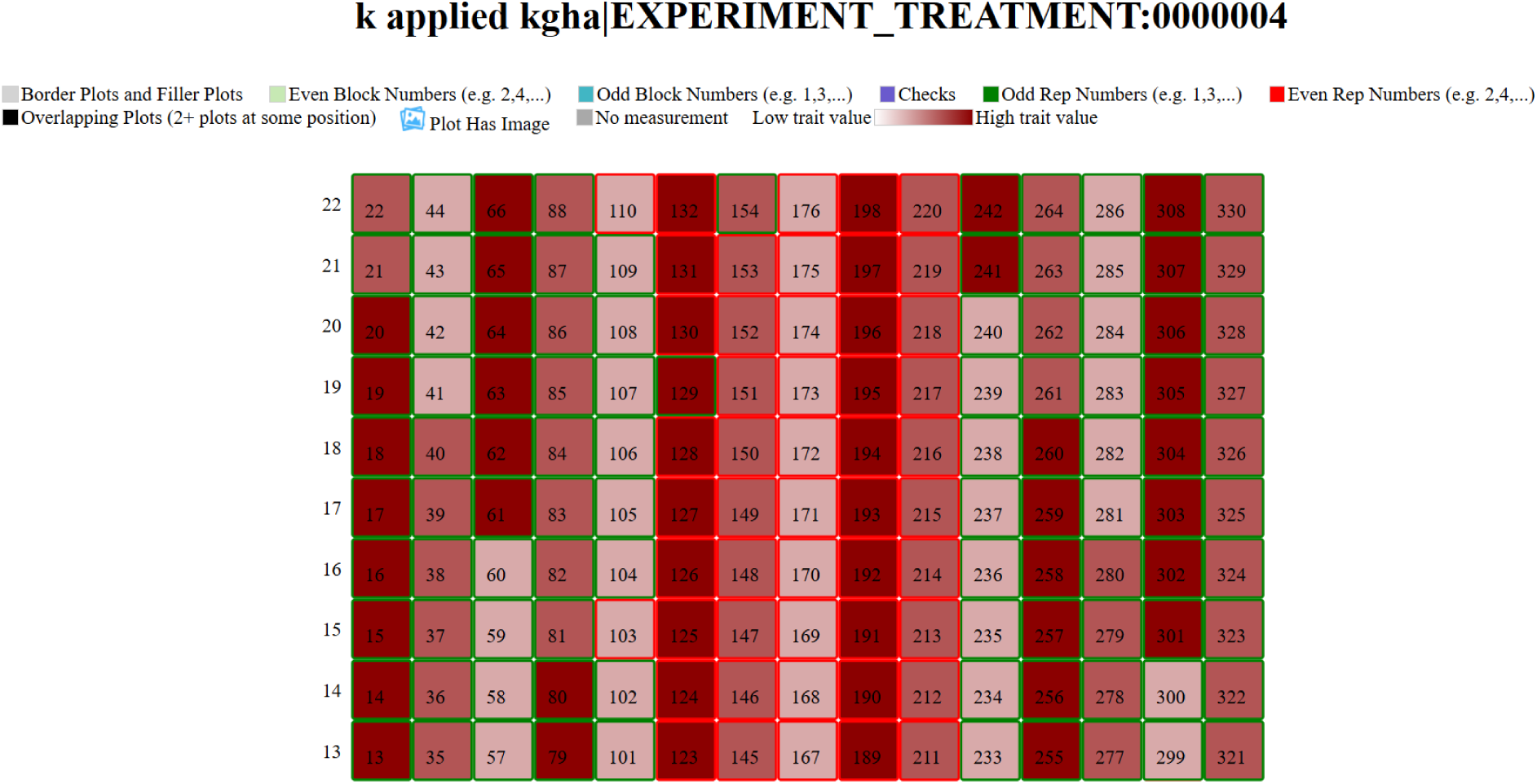
a sample of the application of treatments to plots in a trial. The field map viewer function of Breedbase enables detailed inspection of trait and treatment data across the entire field. By modeling treatments like phenotypic data, experimental designs can be created more flexibly and visualized as a heatmap.

To illustrate the difference in modeling power between Breedbase 1.0 and Breedbase 2.0, we will examine two studies and describe how both versions of Breedbase would handle them.

Let us first consider (López-Bellido et al. 2005). In this study, López-Bellido et al. investigated the timing of nitrogen fertilizer application on Mediterranean wheat, testing whether the timing of the same quantity of fertilizer throughout the wheat lifecycle had a differential effect on yield. They used a complete randomized block design with nine treatments, one control with no nitrogen applied, and four replications for a total of 40 plots. The sum total of nitrogen in each treatment plot was kept constant at 150 kg N ha^-1^, but the timing of nitrogen application was changed between the sowing, tillering, and stem elongation stages across treatments: 150-0-0, 100-50-0, 100-0-50, 75-75-0, 75-0-75, 50-100-0, 50-50-50, 0-150-0, and 0-75-75 kg N ha^−1^. Nitrogen was applied in the form of urea during sowing and ammonium nitrate otherwise.

To model this trial in Breedbase 1.0, we would have had to make a decision between capturing all individual treatment applications or only storing data for the unique treatment combinations described in the paper. Retaining only the unique treatment combinations would have necessitated specifying ten columns in the trial design file for the nine treatments and the control. Users may have chosen to label columns ranging from `0_0_0` to `50_50_50` to capture all possible values and specify a 0 or 1 to place a plot within that treatment group. Had the user wished to store additional data, such as the dates of application and the more detailed description of the nitrogen sources applied at different times, they would have needed to individually and manually specify these data in the descriptions of each treatment after the trial was stored in the database. The other option, to store data related to each individual treatment application, requires specifying 13 columns in the trial design file to account for the five possible sowing-time nitrogen applications, five possible tillering nitrogen applications, and the three possible stem elongation nitrogen applications. They could have chosen to use column names such as `N_sowing_50` or `N_tillering_75` to signify the treatments. As before, once the trial design had been stored, the user would then have needed to manually edit the virtual trials for each of the treatment applications to store additional descriptive data. These setups would have been sufficient to store all the information related to the trial in López-Bellido et al. 2005, but in the context of larger breeding programs, there was no way to search for all trials with nitrogen fertilizer applications and compare their methods programmatically.

Now let us model the same experiment using Breedbase 2.0. Instead of making individual treatments for each unique treatment combination, we can create just two treatments in the treatment ontology - `N_applied_NH4NO3_kgha` and `N_applied_urea_kgha` – to capture the two methods of nitrogen application used in this study: ammonium nitrate and urea, measured as kg ha^-1^. In the treatment designer, we give it a detailed description and specify it as a continuous variable with repetitive applications allowed. We could even group these treatments under a larger treatment group called `N_applied`, which could itself be part of `Fertilizer_treatments`. These treatments would be assigned an ID in the `EXPERIMENT_TREATMENT` ontology, the dictionary of treatment terms used in a Breedbase instance. This organization into a treatment ontology allows us to search for treatment applications, trials, and stocks based on the types of treatments that were applied to them. After creating our treatments, we design a spreadsheet where each row is a plot and the columns are the two treatments we devised. This spreadsheet format should be familiar to existing Breedbase users because it is identical to the phenotyping spreadsheet format. Each entry would be the value of the fertilizer applied followed by a timestamp, which tells us in what stage of the experiment the treatment was applied. For example, a plot with the sowing-tillering-stem elongation treatment of 100-0-50 kg N ha^-1^ would have spreadsheet entries as in Table 1.

**Table 1:** experimental treatment designs reimagined. Treatments can be applied to individual stocks at any stock level, and they may come with timestamps to signify exact time of application.

| **observationunit_name** | **N_applied_NH4NO3_kgha \|**<br>**EXPERIMENT_TREATMENT:0000001** | **N_applied_urea_kgha \|**<br>**EXPERIMENT_TREATMENT:0000002** |
| --- | --- | --- |
| plot_1 | 0,2026-11-01 00:00:00-0000 | 100,2026-11-01 00:00:00-0000 |
| plot_1 | 0,2027-01-15 00:00:00-0000 | 0,2027-01-15 00:00:00-0000 |
| plot_1 | 50,2027-02-28 00:00:00-0000 | 0,2027-02-28 00:00:00-0000 |

This scheme produces the expected outcome of 100 kg N (urea) ha^-1^ during sowing in November, no nitrogen during tillering in January, and 50 kg N (ammonium nitrate) ha^-1^ during stem elongation in late February, and it does so with less effort and greater clarity.

Alternatively, storing time data could be done by composing a treatment with a time ontology trait into a single treatment reflective of both. This approach is commonly used by Breedbase users when creating traits that capture the same physiological data but at specific times in development or in the agricultural cycle. Because treatments are now implemented in much the same way as phenotypic traits, they may also be composed with entries in the time, unit, method, and plant tissue ontologies to form rigorous, fine-grained definitions of the treatment values. The flexibility of treatments in Breedbase 2.0 ensures users have the choice to store treatment data reproducibly using either approach.

Our second study is (López-Bellido et al. 1998). In this study, López-Bellido et al. tested three treatment types in a split-plot design: tillage, crop rotation, and nitrogen fertilizer. Plots were assigned to be tilled conventionally (CT) or not tilled (NT). Plots were split into subplots for each two-year crop rotation method: wheat-sunflower (WS), wheat-chickpea (WCP), wheat-faba bean (WFB), wheat-wheat (WW), and wheat-fallow (WF). Subplots were split into sub-subplots with nitrogen application levels of 0, 50, 100, and 150 kg N ha^-1^.

In Breedbase 1.0, modeling these data using the split-plot design in the trial design workflow is only possible by creating treatment names for all unique combinations of treatments (20 for nitrogen applied and crop rotation) and applying a tillage value to the plots afterwards. This is sufficient to capture the range of treatments applied to each plot, but it necessitates the laborious creation of 20 individual treatment names. Data modeling becomes difficult because the user must convert this overencumbering one-hot encoding to the original treatment values to be useful for downstream analyses.

Breedbase 2.0 supports subplots but not sub-subplots. Nonetheless, the new treatment implementation makes storing these data simple in comparison. We can reuse the `N_applied_NH4NO3_kgha` treatment from the previous example and create additional treatments for `tillage` and `crop_rotation`. These treatments can be defined as categorical and restricted to a predefined set of valid values. Using the split-plot design in the trial design workflow, we can specify the treatments and levels of those treatments that must be present in each plot. In our case, a subplot will be created for each unique combination of `crop_rotation` and `N_applied_NH4NO3_kgha` to create 20 subplots per plot. The parent plots can then be specified a `tillage` value using the spreadsheet method as before. While this does not capture the full depth of the original sub-subplot structure, this can be easily accounted for in downstream statistical analyses because observational units can be blocked by treatment values shared in common, which reproduces the sub-subplot structure of the original experiment. This trial and others later can utilize the same treatments, enabling users to search for trials based on applied treatments and streamlining the process of designing and storing trials in Breedbase.

## Repetitive Measurements

In many modern breeding programs - including horticultural crops like strawberries, forages like alfalfa (Thapa et al. 2024), or other perennial crops - phenotypic data collection happens throughout the season rather than just once at harvest at the end of the trial (or experiment) as is often the case with cereal crops (Hale et al. 2025). Breeders in these programs frequently record observations like plant height, disease scores multiple or yield times over the duration of an experiment. In Breedbase 1.0, implementing time series measurements required generating so-called post-composed terms between the trait variable of interest and a time ontology term (Menda et al. 2025), as well as potentially other ontologies, such as sample preparation ontologies. Post-composing remains the preferred method for managing time series data for many use cases, particularly when data will only be compared within the post-composed categories. In certain conditions, however, it is more convenient to simply take a large number of time-based measurements, and store the phenotypic values together with a timestamp and other metadata. Using the timestamp information, phenotypic data can be queried as time series data, or queried in specific data ranges. To address this use case, we added better support for repetitive measurements to Breedbase 2.0. Changes to the system involved how repetitive traits are managed and how repetitive phenotypic data are collected, displayed and queried.

### Trait Variable Attributes

In Breedbase, traits are organized into ontologies, which represent hierarchical graphs where a root node branches into subcategories via `is_a` relationships. While traits themselves represent general concepts (e.g., “plant height”), actual measurement data requires the creation of trait variables. These variables ensure reproducibility by linking to three specific entities: a trait (`variable_of`), a method (`method_of`), and a scale (`scale_of`). Furthermore, trait variables support attributes that define data types (e.g., categorical, numerical, or timestamp), category names, and value constraints.

Repetitive data can be reported in different ways. For example, when querying a database for a trait with repetitive values, all the measured values could be reported to the user. However, sometimes the user is more interested in the first value, the last value, the sum of all the values, and most often in the average value. Therefore, when querying phenotypic data, the reporting mode has to be specified, with the possible values “all”, “first value”, “last value”, “sum”, and “average”.

To accommodate the different flavors of repetitive data, a new attribute called `repeat_type` was introduced, allowing traits to be categorized as `single`, `multiple`, or `time_series`. A `time_series` trait is measured many times and in association with its other measurements throughout a trial or experiment. For instance, plant height could be measured several times to generate a growth curve. Because aggregating the `time_series` data via simple averages is often statistically inappropriate for downstream applications such as mixed models or Genome-Wide Association Studies (GWAS) (Tecle et al. n.d.), a different repeat type should be selected for these analyses.

A typical use case for the `multiple` repeat type is a trait that is measured over long periods of time but does not represent a time series. Good examples are yield-related traits in perennial crops such as strawberry. Strawberries have a long harvesting period during which harvesting occurs continuously, and the yield is measured many times. The data represent the same yield trait, but are associated with different harvesting times; with the `multiple` trait, the system allows querying the difference in yield early compared to late in the harvesting season. The default reporting mode for `multiple` traits is “average”.

For example, consider the constraints of a strawberry breeder: strawberries may have multiple harvests over a single growing season (Holmes 2024). Suppose the strawberry breeder is evaluating accessions at the plot level to determine which elite varieties will progress to the next step of the breeding pipeline. If only one measurement is allowed for fruit yield using the total weight method (`CO_372:0000004`), the breeder will be forced to take multiple measurements over the course of the season, sum them, and enter them into their breedbase instance only after the harvest season is over. They are forced to return to manual data management on local spreadsheets, which reintroduces some of the problems Breedbase aims to solve. In Breedbase 2.0, the breeder may instead store yield as a repeated measurement of type `multiple`, enabling data management in the cloud on Breedbase. By pairing their trait measurements with timestamps, they can ascertain yield over the course of the harvest season in addition to plot-level comparisons of accessions. The structure of the data is conserved, and the breeder has better documentation for their advancement decisions.

When multiple or time_series are selected, the measurement will require a time stamp along with the measured value, which will be used to distinguish the measurements from previously taken measurements. It also allows the measurement to be queried by time range.

### Repetitive measurements user interface

We developed a repetitive measurements data viewer (see the as a visual interface to explore the quantitative repeated phenotyping measurements (figure 2). The technical architecture of this viewer emphasizes cross-platform compatibility and responsiveness.

- Visualization: the interface utilizes the D3.js (https://d3js.org/) javascript library, a powerful tool for creating interactive web-based data visualizations using hypertext markup language (HTML), cascading style sheets(CSS), and Scalable vector graphics (SVG).
- User Interface (UI): The UI was built using the Bootstrap framework (https://getbootstrap.com/), providing a responsive design that adapts to the screen of any device researchers may use both in the field and the office.
- Data Handling: The data viewer is specifically designed to handle multiple or time-series data points for traits like yield and plant height, allowing users to visually track quantitative changes across different dates throughout the trial’s duration (figure 3).

**Figure 2:**
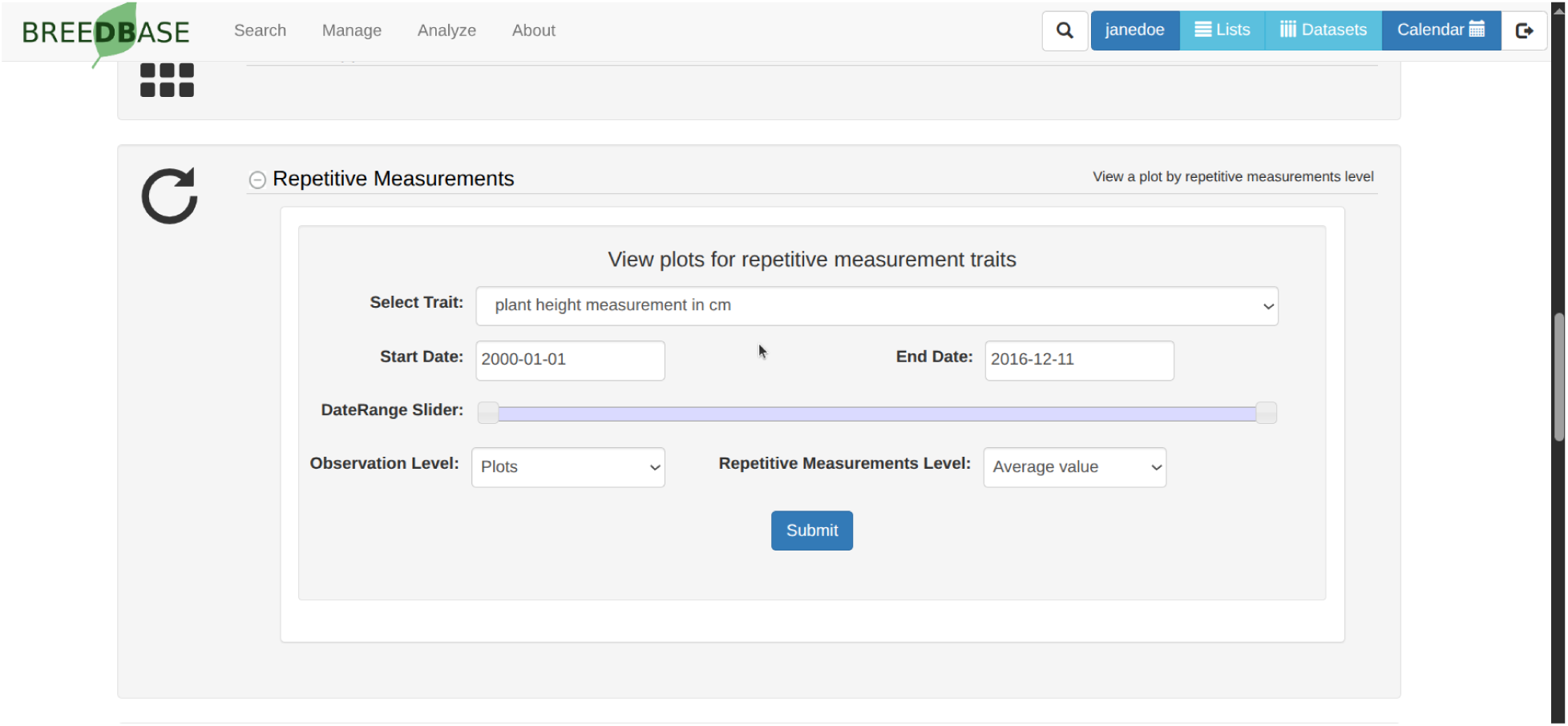
Repetitive measurement section showing different data handling features on the breedbase

**Figure 3:**
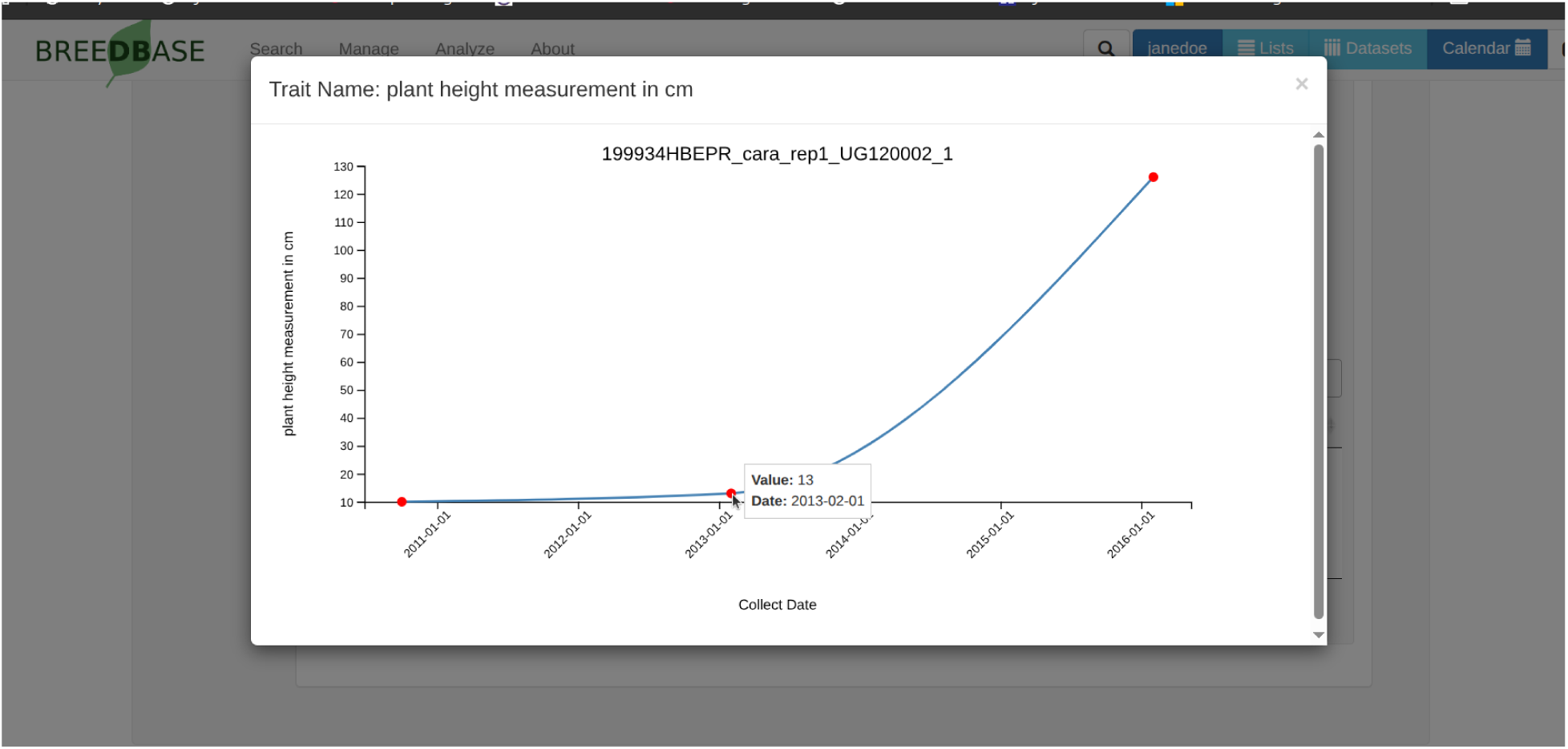
Visualization of time series data, showing values and dates for the repetitive trait

Additionally, we integrated the repetitive measurement sections into the data download system to provide researchers with flexible aggregation options. When exporting phenotypic data, users can choose different options (first, last, average, sum) of measurements, or alternatively retrieve all individual observations with timestamps included. To accommodate diverse analytical workflows, the tool allows for the inclusion or exclusion of temporal timestamps and supports precise filtering based on specific date ranges or numerical value thresholds

By transitioning from a terminal-harvest model to repetitive measurement model, Breedbase 2.0 allows for a more nuanced understanding of plant development. This dedicated section simplifies the process of identifying temporal patterns, such as the rate of biomass accumulation or the speed of disease onset, which are critical for selection in perennial and multi-harvest breeding programs.

## Phenotype Quality Control

Ensuring high data quality (Regt 2020) is a major concern in breeding programs because the reliability of downstream analyses and selection decisions depends directly on the accuracy and consistency of the collected observations. Within the Breedbase digital ecosystem, data quality checks are incorporated at multiple stages of the data acquisition workflow. For example, the use of barcodes in the field to identify plots, plants, and other experimental units can substantially reduce material identification errors. Similarly, data type restrictions in Field Book, including predefined categorical values and limits for numerical traits, can reduce data entry errors during field collection. Additional validation is performed when phenotypic data are uploaded to Breedbase. Nevertheless, erroneous or experimentally unreliable observations may still enter the database as a result of measurement errors, field conditions, transcription problems, or compromised experimental units. Consequently, additional tools are required to identify, document, and, when appropriate, exclude problematic observations from downstream processing and analyses.

In Breedbase 2.0, a phenotype quality-control (QC) workflow was implemented to help breeding teams identify and review potentially problematic field observations before downstream analysis. The interactive interface was developed in JavaScript to provide rapid visual feedback and a streamlined review-and-apply workflow. Rather than automatically deleting observations identified as unusual, the QC process supports user-guided evaluation of potentially problematic data. The original phenotypic records remain preserved, while observations identified through the QC workflow can be flagged and subsequently excluded from compatible data retrieval and analysis procedures. This approach maintains the integrity of the originally collected data while preserving the provenance of QC decisions.

An important distinction in phenotype QC is the difference between dataset-level and database-level quality control. In dataset-level QC, outliers are identified only within the specific dataset in which the QC procedure was performed. Consequently, if another user creates a new dataset containing the same trials and phenotypic observations, the previously detected outliers are not automatically recognized as such, even when the new dataset contains the same experimental records. The QC procedure must therefore be repeated for each independently created dataset (figure 4). In contrast, database-level QC records the outlier status at the individual phenotype observation level. Once an observation has been reviewed and validated as an outlier, this QC status persists independently of the dataset in which the observation was originally evaluated. Any subsequent dataset containing the same phenotypic observation can therefore recognize and, when requested, exclude the previously identified outlier. The Breedbase phenotype QC tool described here follows this database-level approach, allowing validated outlier classifications to be retained and consistently applied across datasets and downstream analyses.

**Figure 4:**
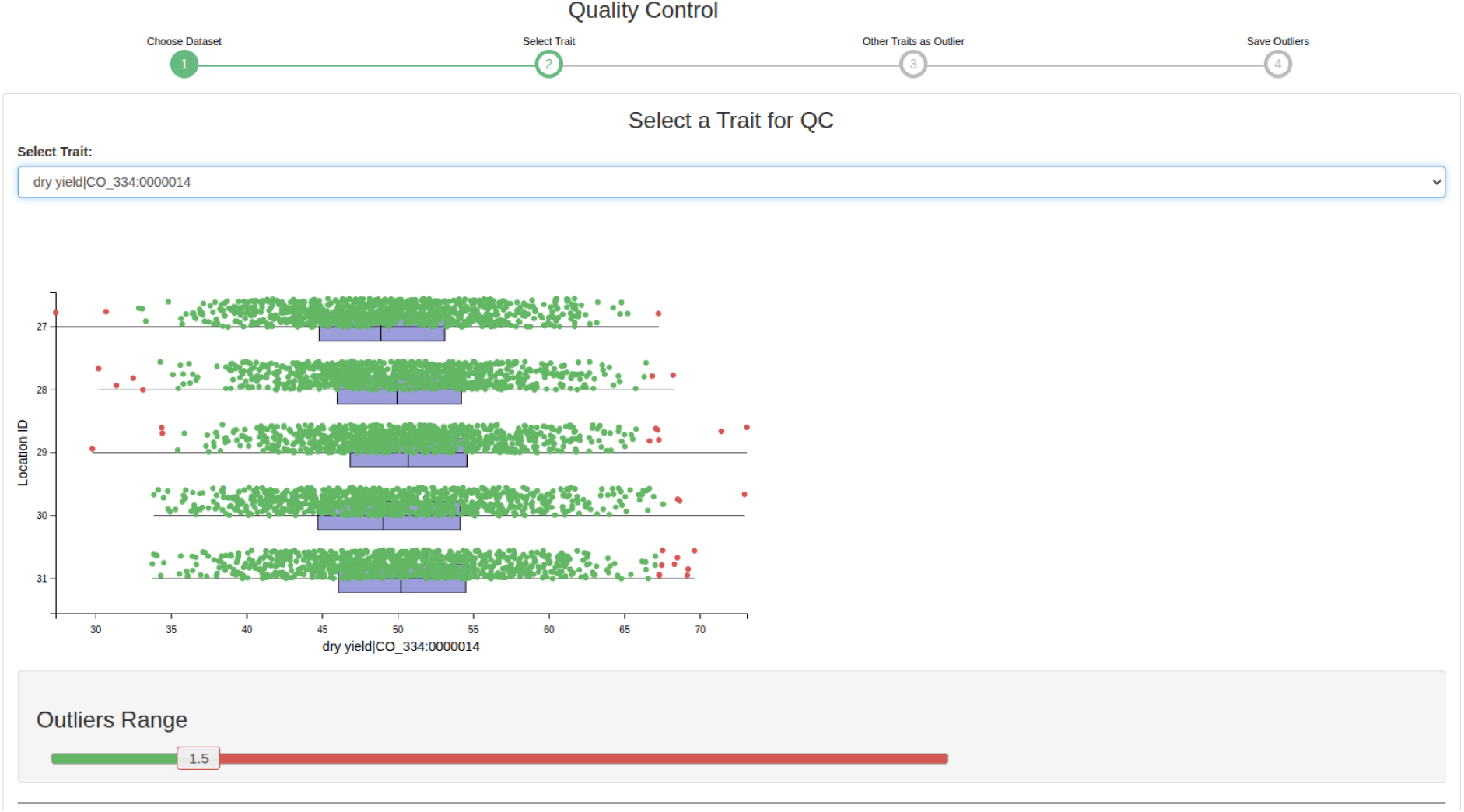
The quality control feature tool with a sample trait and distributions. Red dots indicate measurements that have been marked as outliers. The outlier range can be adjusted with the slider to change the IQR cutoff, with the display changing outliers in real time.

This persistent phenotype-level QC status enables users to work with validated observations through validated trait datasets. In practice, downstream analyses, including mixed-model procedures for Best Linear Unbiased Prediction or Estimation (BLUP/BLUE), can be configured to automatically exclude phenotype observations previously validated as outliers. This reduces the risk that known data-entry errors, anomalous measurements, or observations from compromised experimental units are inadvertently included in routine analyses. At the same time, additional dataset-level filtering can still be applied in a user- and analysis-specific manner when researchers require criteria that are relevant only to a particular analysis. Thus, database-level QC provides persistent and reusable phenotype-level outlier identification, whereas dataset-level QC remains temporary and specific to the dataset in which the filtering criteria were applied.

A key component of the database-level QC workflow is statistical outlier detection using an interquartile range (IQR)-based approach (figure 6). Users select a dataset and a trait of interest, and the distribution of phenotypic observations is displayed across the experimental locations represented in the dataset. Potential outliers are visually distinguished from the remaining observations. The IQR multiplier used to define the outlier range can be interactively adjusted through a slider, allowing users to evaluate different levels of stringency according to the characteristics of a trait, trial, or environment. This interactive approach is important because statistical outliers do not necessarily represent erroneous observations; they may reflect measurement problems, localized environmental effects, or biologically meaningful extremes. Therefore, identified observations are presented for review before their outlier status is persistently recorded in the database.

The QC workflow also supports trait-specific validity constraints through predefined minimum and maximum values. This functionality is particularly useful for numerical, ordinal, or numerically encoded scoring traits with established measurement ranges. For example, plant disease severity may be evaluated using a defined 1–9 scale, for which observations outside the expected range generally indicate data entry errors, inconsistent scoring conventions, or transcription problems. Trait-level limits provide an additional standardized validation layer for phenotype records stored in Breedbase. These constraints differ from contextual statistical outlier detection because they identify observations that violate the defined measurement properties of a trait rather than observations that are statistically unusual within a particular set of experiments.

The workflow further enables cross-trait, or gating, QC based on a primary trait. In field experiments, the reliability of one phenotype may depend on the condition of the experimental unit as described by another measurement. For example, a plot with an insufficient number of viable or harvestable plants may provide an unreliable estimate of plot-level yield or other secondary characteristics. After identifying an experimental unit as problematic based on a primary trait, users can select additional traits for which observations from the same plot should also be flagged as outliers. Thus, low stand count can be used as evidence that selected yield or secondary trait observations from the same plot are not sufficiently representative for downstream analysis.

This cross-trait functionality differs from both fixed trait validation limits and conventional statistical outlier detection. The secondary phenotype value itself may fall within the expected biological range and may not be statistically extreme; however, the experimental condition under which it was measured may compromise its interpretation. The gating workflow therefore allows QC decisions to reflect relationships among traits and the overall quality of an experimental unit. For large multi-trait trials, this approach reduces the need to manually identify the same compromised plots across multiple traits and helps ensure that consistent exclusion criteria are applied. At the same time, users explicitly select the secondary traits affected by the primary QC decision, avoiding the assumption that every measurement collected from a flagged plot should automatically be excluded.

Breedbase also records phenotype validation information to support the traceability of the QC process. Trials and traits that have undergone QC can be retrieved together with information describing the experiment, location, evaluated trait, and the user responsible for the validation. This enables breeding programs to distinguish phenotypic data that have undergone formal review from observations that have not yet been evaluated. The resulting validation history provides provenance for QC activities and is particularly valuable in collaborative breeding environments, where phenotype collection, curation, validation, and analysis may be performed by different members of a research team.

Validated phenotype data can also be tracked across research locations through a geographical interface. Trials that have undergone QC are associated with their experimental locations and can be displayed on an interactive map, providing an overview of phenotype validation activities across the experimental network. For multi-location and multi-environment breeding programs, this visualization enables researchers and data managers to identify locations and trials for which QC has been completed and to recognize experimental environments that may still require review. The map therefore provides both a data provenance resource and an operational overview of phenotype validation across research sites (figure 5).

**Figure 5:**
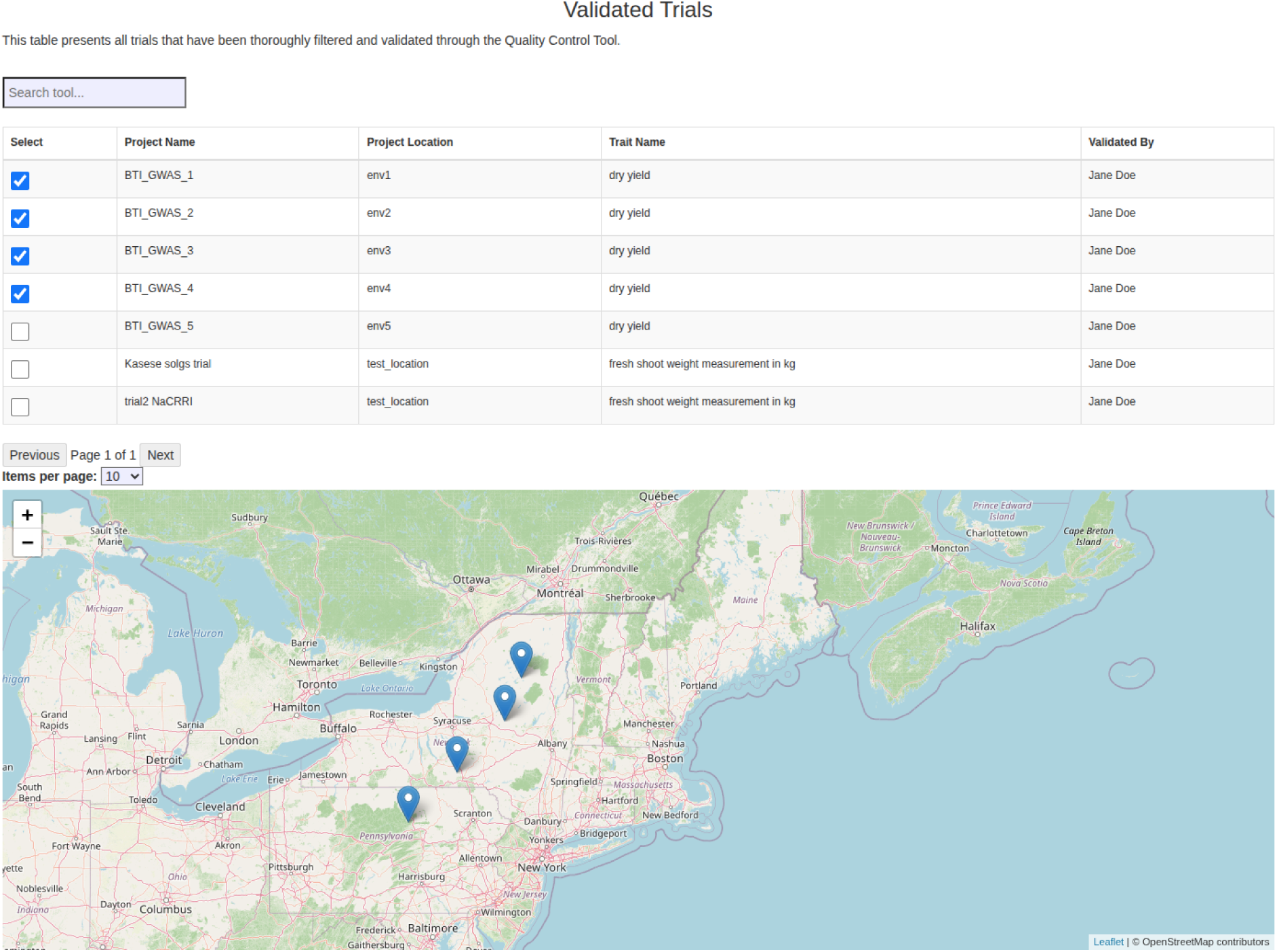
Screen showing the validated trials, including basic information about the trials in tabular form, and a map of trial locations.

**Figure 6:**
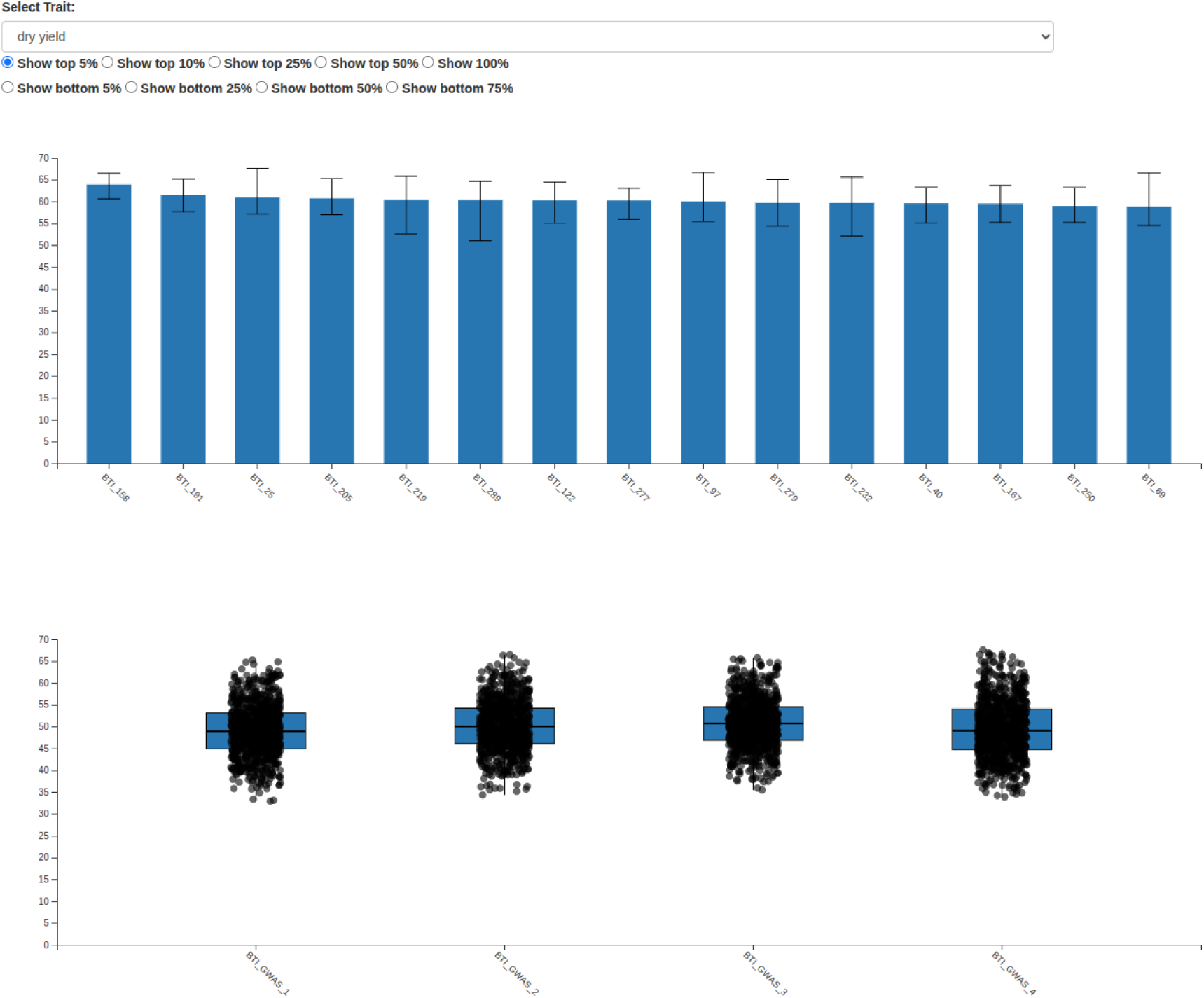
Graphical view of performance for a given trait (dry yield) for the top 5% of lines, as well as distribution of trait scores in the validated trials.

In addition to identifying and recording potentially problematic observations, the QC environment provides statistical and graphical summaries that support phenotype review. Accession-level descriptive statistics can be calculated for a selected trait, including the number of trials and observations represented, mean, standard deviation, minimum, and maximum values. These summaries can also be exported for further evaluation. Bar plots allow users to visualize and rank selected proportions of the highest- or lowest-performing accessions, while boxplots provide a representation of phenotypic distributions and variability across accessions or experimental groups. Together, the statistical summaries, bar plots, and boxplots provide additional context for identifying unexpected data patterns and evaluating phenotype consistency before subsequent genetic or breeding analyses.

## Intercropping Support

The trial structure for Breedbase 1.0 was originally designed to support a monocrop breeding system. In this system, a crop of a single species is grown in monoculture in the trial and a single accession of this one crop is grown within each plot of the trial (Vandermeer 1989). Monocropping systems have become more widespread over the last few decades due to the development of elite modern varieties from the “green revolution” (Evenson and Gollin 2003) that have higher yields and have allowed food production around the globe to increase (Conway 1998). The data structures used to store trial data within Breedbase 1.0 were designed with this system in mind.

Intercropping is a multiple cropping system where two or more crops are grown in the same space at the same time, creating inter-crop competition or synergy during some or all of the crop growth (Andrews and Kassam 1976). It is likely one of the earliest and most widely performed agricultural practices (Plucknett and Smith 1986) and still remains widely employed throughout the world, particularly in small-scale farms (Brooker et al. 2015). Intercropping has gained some renewed interest as a way to offset some of the detrimental environmental impacts of monocrop systems, due to improved pest control and more efficient uses of water and nutrients (Tilman et al. 2002; Moore et al. 2023).

In order to support intercropping in a data management system, there are three main features that need to be included:

1. The database needs to be able to store the accession records from multiple crops and clearly specify which crop or species the accession belongs to
2. The plots within the trial need to be able to include more than one accession record
3. The database needs to support more than one trait ontology.

Breedbase 2.0 has been updated to address these needs in order to better support intercropping systems.

### Storing accessions of multiple crops

The ability to create and store accession records for multiple crops is already supported by Breedbase. Each accession record has a required organism property which stores the genus and species name of the crop the accession belongs to. By default, a Breedbase instance contains the entire Genbank plant taxonomy, but this can be extended or adapted as needed. This property is displayed prominently on the detail page for each accession record and is included in the table of search results when looking up existing records in the database. The user can also filter accession records by organism, allowing them to find accession records from a specific crop. All of these features allow Breedbase to easily store accession records from multiple crops used by an intercropping system.

### Storing multiple accessions in a single plot

The data structure used to store a trial in Breedbase did not allow more than one accession to be associated with a plot. This limitation prevented any intercropping trials, which have more than one accession grown within a single plot, to be stored in Breedbase. A plot in a trial (stored in the `stock` table with a type of *plot*) and the accession grown in the plot (stored in the `stock` table with a type of *accession*) are directly linked to each other with an entry in the `stock_relationship` table using a *plot_of* relationship type, where the id of the plot is set as the `subject_id` and the id of the accession is set as the `object_id` (Figure 7).

**Figure 7:**
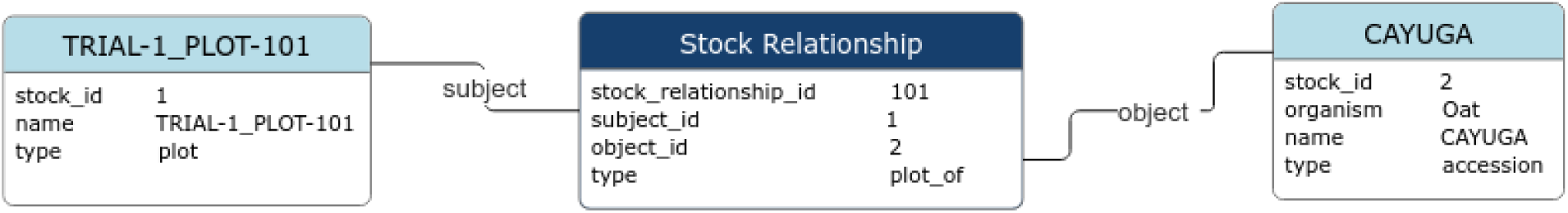
A database relationship diagram showing the connection between a plot and an accession using the *plot_of* relationship type.

There are numerous functions within the existing Breedbase codebase that expect just a single *plot_of* relationship between a plot and the accession grown in the plot. Therefore, a new stock relationship type was created to store additional associations between a plot and any intercropped accessions grown within it. This *intercrop_plot_of* relationship allows for more entries in the `stock_relationship` table between a plot and one or more additional accessions grown in the plot, also storing the id of the plot as the `subject_id` and the id of the intercropped accession as the `object_id` (Figure 8).

**Figure 8:**
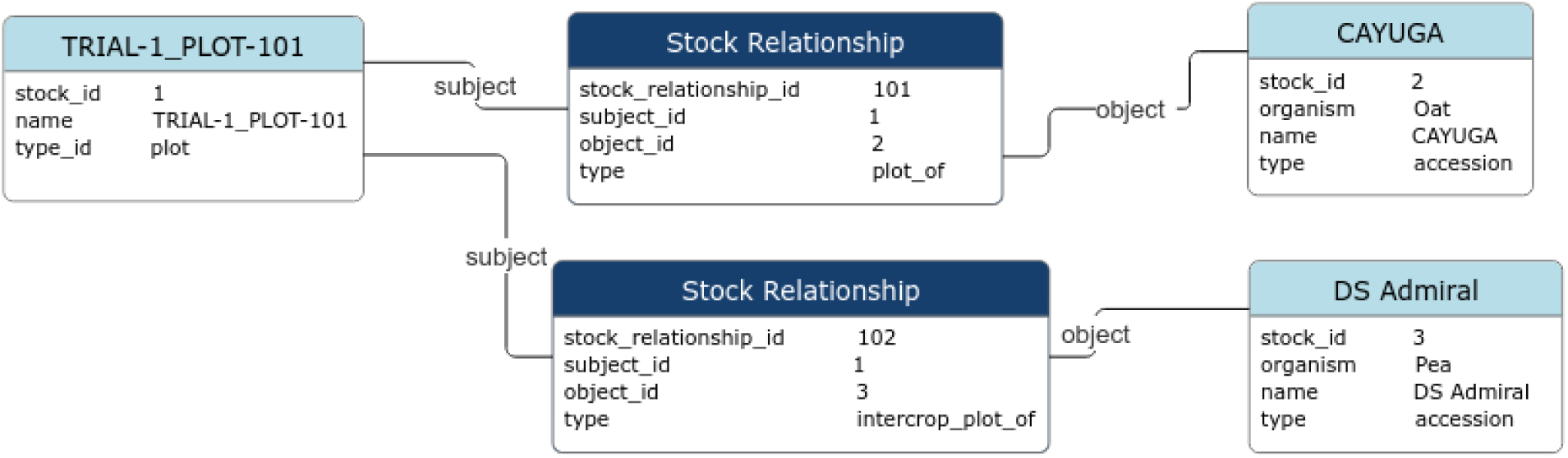
A database relationship diagram showing the new *intercrop_plot_of* relationship type that can link additional accessions to a plot.

Changes were made to the trial upload process and user interface of the website in order to allow users to create these new relationships when storing a trial and to display the intercropped accessions of a created trial. A user can create an intercropped trial through a file upload. Generating intercropped trials though the interactive trial design tool on the website is not yet supported. The file contains 1 row per plot and contains metadata about the trial (such as the trial name, breeding program, location, and planting and harvest dates) and metadata about the plot (such as the plot number, the block and replicate numbers, and the row and col position of the plot). The ‘accession_name’ column allows the user to specify the name of the accession that is grown in each plot. The accession entry associated with this name is then linked to the plot using the *plot_of* stock relationship when the trial is created. Recent changes have added an optional ‘intercrop_accession_name’ column to the trial upload file, which can include additional intercrop accession names for each plot. This column can include zero, one, or more comma-separated accession names which will be linked to the plot using the *intercrop_plot_of* stock relationship.

The trial detail page was updated to display intercrop accessions if they have been added to the trial. The Experimental Design section of the detail page includes a table of all of the Accessions that are sampled in a trial. It has been updated to include any intercrop accessions that have been added as well as the species name of each accession to easily distinguish between different crops (Figure 9). The plots table in the Experimental Design section, which lists all of the plots in a trial along with their associated accession, has also been updated to include any intercrop accessions that have been added (Figure 10). The spatial layout display on the website, as well as the download, now includes the intercrop accessions as well (Figure 11).

**Figure 9:**
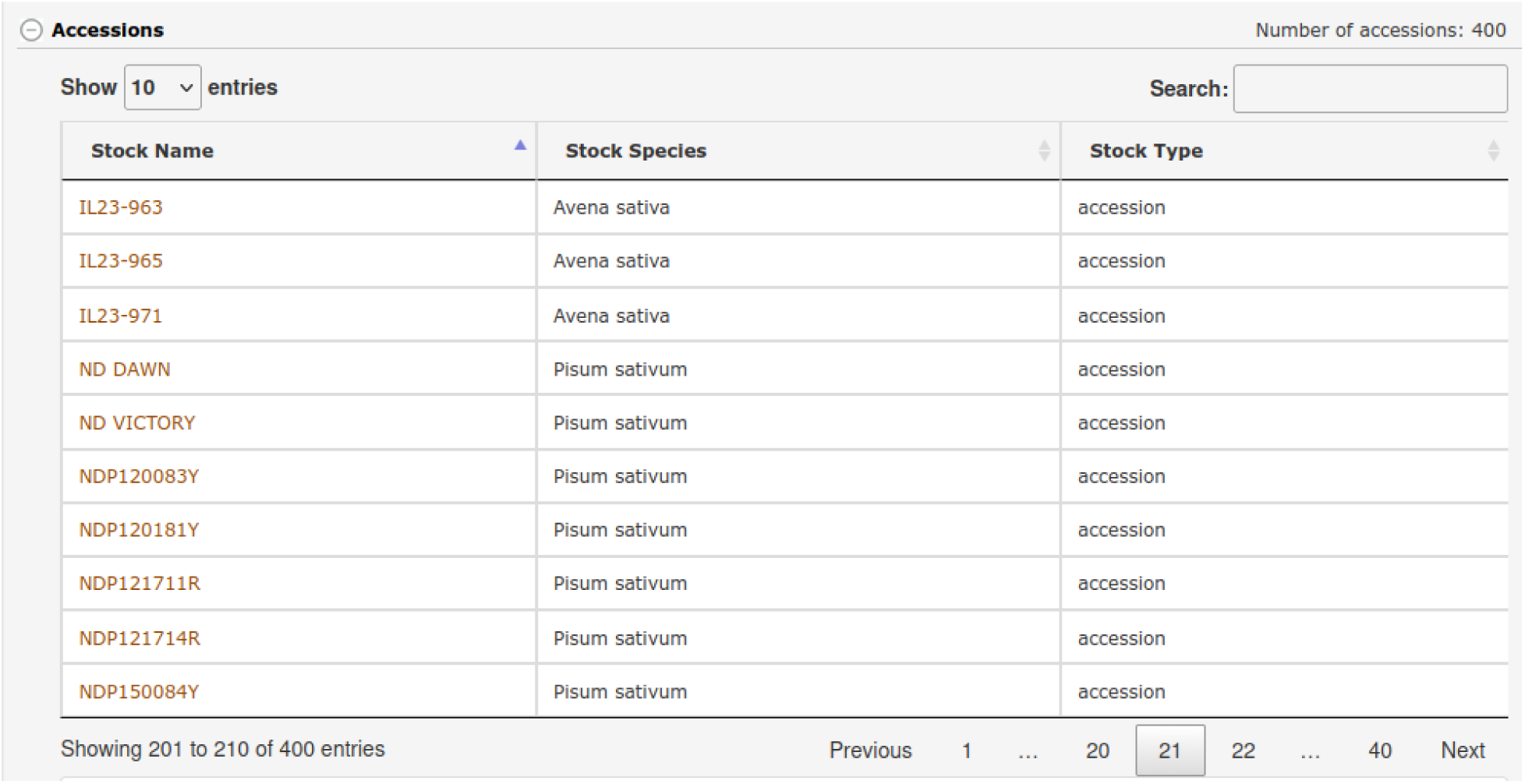
The Accessions table on the trial detail page lists all of the accessions that were included in a trial, now including any intercrop accessions along with the species name for each one.

**Figure 10:**
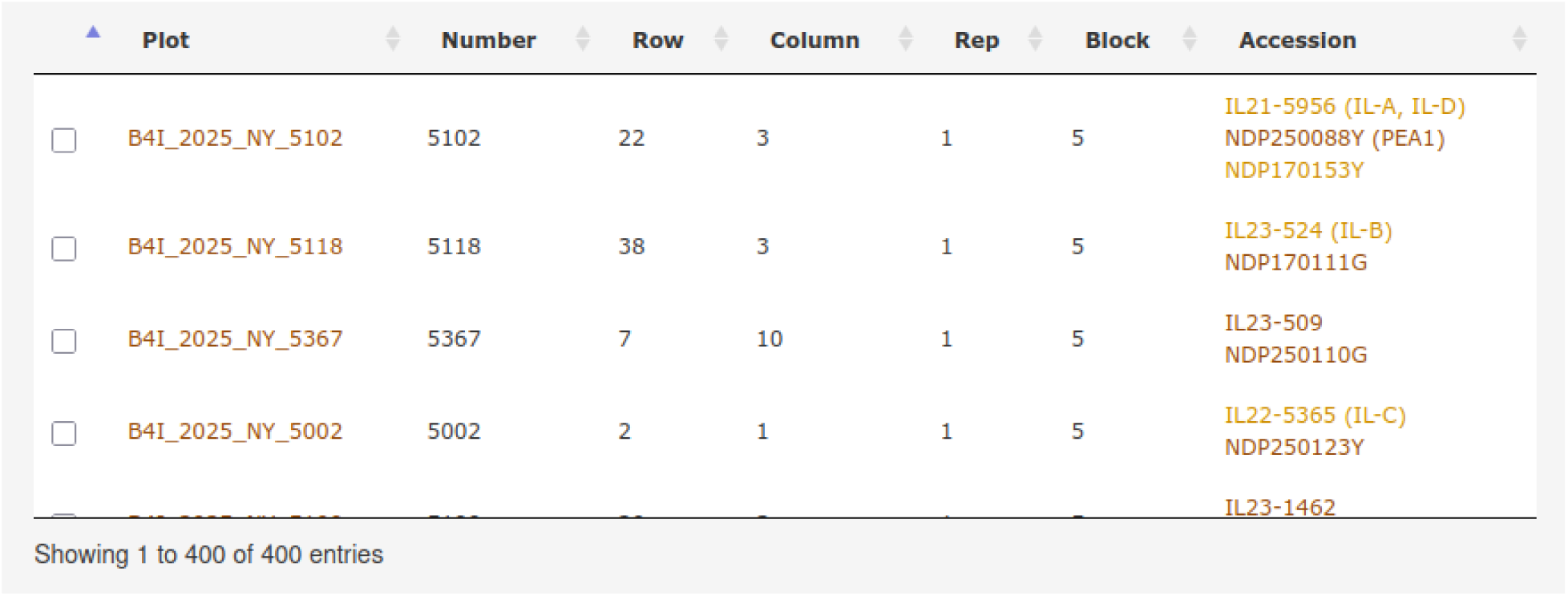
The Plots table on the trial detail page now lists all accessions associated with each plot.

**Figure 11:**
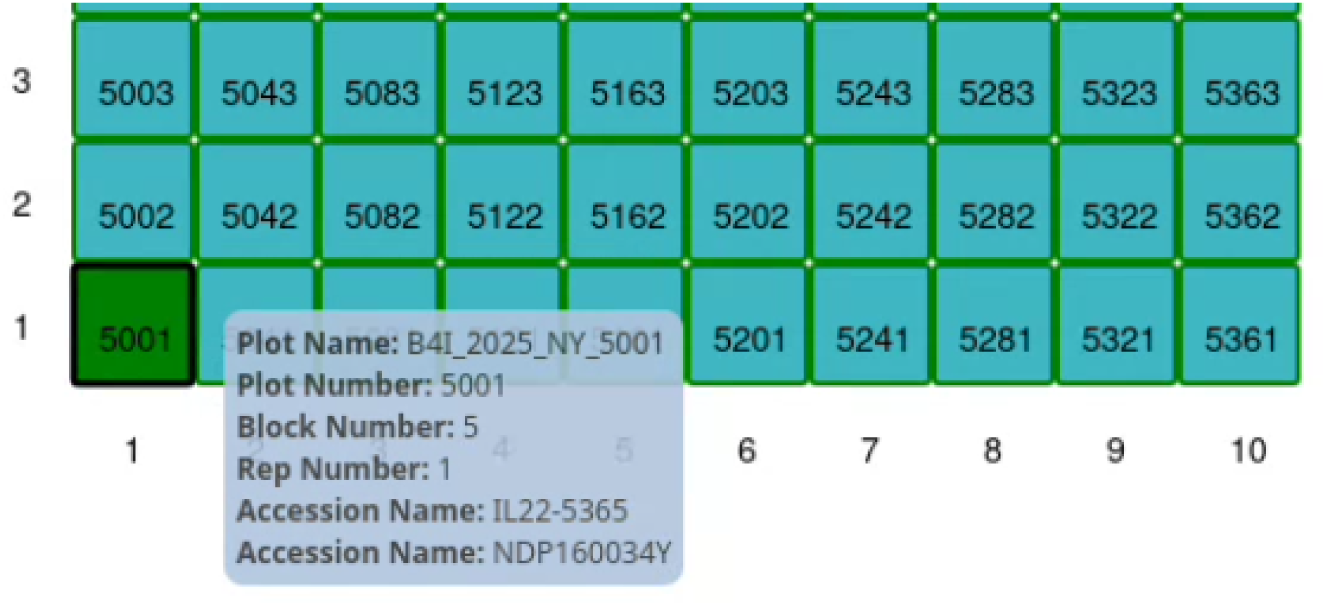
The plot tooltip (displayed when hovering over a plot) on the trial layout on the trial detail page now lists all accessions associated with the plot.

Including the intercrop accessions in the phenotype downloads is of critical importance for any downstream analyses using the data stored in the database. The downloads include plot-level metadata along with any trait observations made in each of the plots in the selected trials. The plot-level metadata includes columns for the id and name of the accession in the plot and has been updated to include the id and name of any additional intercrop accessions also in the plot.

The Breedbase Search Wizard is one of the easiest and fastest ways to explore data in the database. The Search Wizard is already capable of displaying which accessions are included in a particular set of trials and/or plots. It has been updated to also include intercropped accessions. The user can easily select a trial that they are interested in and quickly view the list of all accessions that have been added to that trial. The reverse is also possible: the user can select an accession in the Search Wizard and easily find all of the trials that accession has been used in, including its use as an intercrop accession.

Users can also access information about trials programmatically via BrAPI. The `/observationUnits` endpoint can be used to fetch metadata about the plots within a trial. The BrAPI specification defines the `germplasmName` property as the name of the germplasm record (or accession) that is used in a particular plot. As of version 2.0 of the BrAPI specification (Selby et al. 2025), a plot is only allowed to include a single accession entry. Discussions are underway around supporting multiple accessions per plot in a future version of BrAPI. However, until the BrAPI specification is updated, Breedbase will add the names and IDs of intercropped accessions to the `additionalInfo.intercropGermplasm` property of each observation unit (or plot) that contains intercropped accessions.

### Supporting more than one Trait Ontology

Breedbase uses ontologies (Smith et al. 2007), or controlled vocabularies, to store and annotate data in a well-structured format that allows it to be interoperable with other data storage systems (Dumschott et al. 2023). Breedbase has already been able to support the storage of more than one ontology within the database. This allowed the database to include the main trait ontology along with additional ontologies, such as the Gene Ontology (Ashburner et al. 2000) to annotate gene and gene products stored within the database. However, Breedbase was initially designed to operate with just a single trait ontology, limiting its ability to store and use separate ontologies for each crop phenotyped within the database.

With a few minor database query adjustments and ontology configuration changes, Breedbase can now easily store and use more than one trait ontology. This allows curators to maintain separate ontologies for each crop used by a particular Breedbase instance. This, in turn, makes it easier to keep the ontologies on the database in sync with the published trait ontologies, such as those on cropontology.org (Shrestha et al. 2012). Users are also able to easily filter the traits by the ontology, making it easier to find traits for a specific crop (Figure 12).

**Figure 12:**
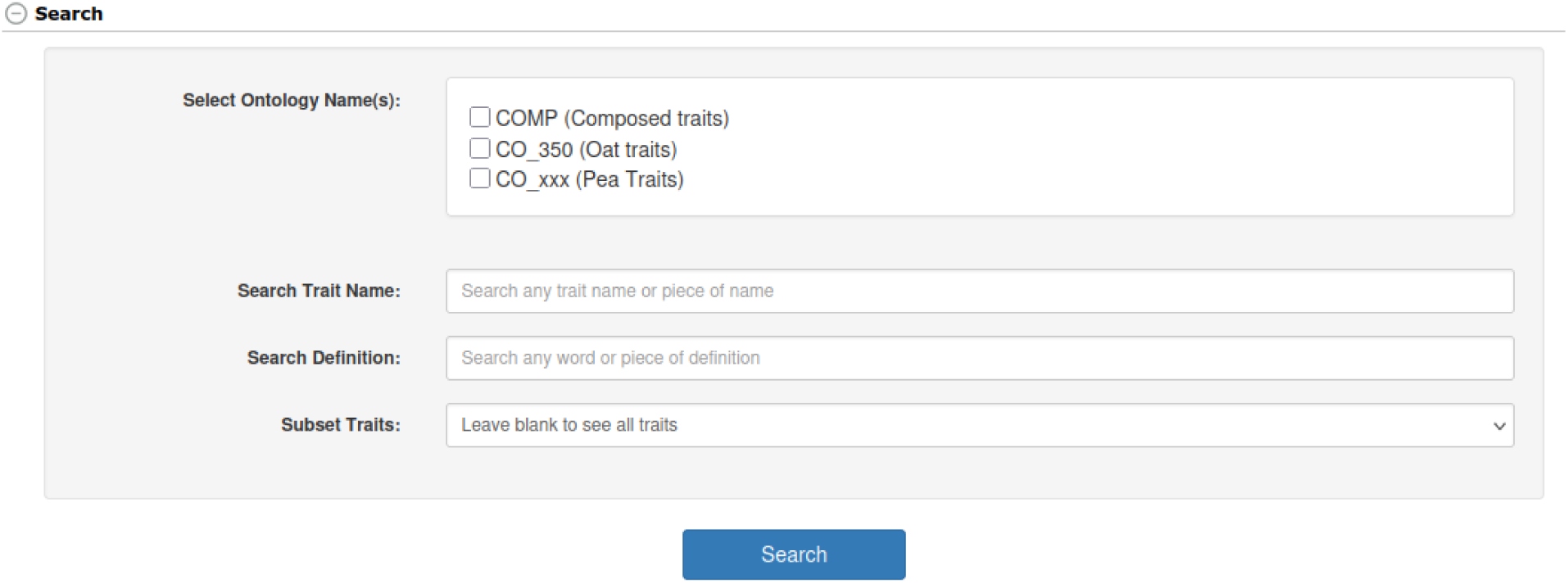
The Trait Search interface allows the user to select which trait ontology to search in, making it easy to find traits for a specific crop when trait ontologies from multiple crops are stored in the database.

Updates to the latest version of Breedbase address the key requirements for supporting intercropping experiments, making it an excellent database system for managing multi-crop trials. The system now supports storing accessions from multiple crops with clear species identification, allows multiple accessions to be associated with individual plots, and enables the management of separate trait ontologies for different crops. These enhancements, integrated with the trial upload process, user interface, data downloads, search functionality, and BrAPI endpoints, demonstrate that Breedbase provides a comprehensive, well-designed solution for researchers conducting intercropping studies who require robust data management capabilities that go beyond traditional monoculture breeding systems.

## Field-level Trial Layout

The Multi-Trial Allocation Tool was developed to support the spatial planning and organization of multiple field experiments within a research station. The tool provides an interactive environment in which users can select an existing location registered in Breedbase and define the corresponding year and season for the planned experiments. Once the location is selected, users can specify the dimensions of the available experimental area in terms of rows and columns, creating a virtual grid (Figure 13) that represents the field or research station layout.

**Figure 13:**
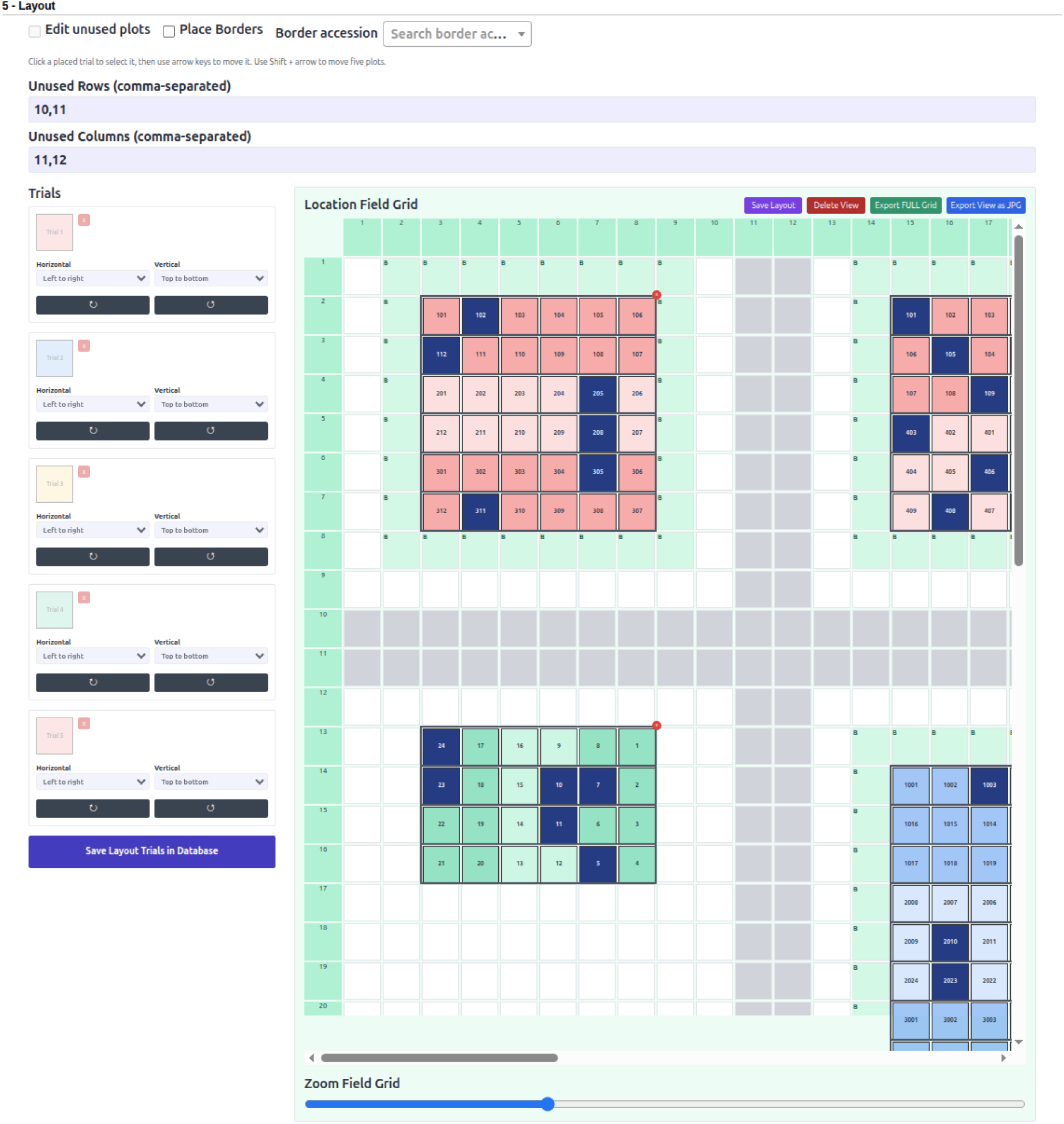
Trial allocation interface showing the drag-and-drop function and field grid layout.

Within this grid, multiple trials can be designed and spatially allocated. Trials can be interactively positioned using a drag-and-drop interface, allowing users to explore different field arrangements and optimize the use of the available space. The tool also provides flexibility to rotate trials, designate plots as unused, and define specific regions within the field. These features are particularly useful when the physical structure of a research station includes irregular areas, restricted zones, roads, infrastructure, or sections that cannot be used for experimentation.

A major objective of the Multi-Trial Allocation Tool is to facilitate coordination among researchers who share the same experimental station. In many research programs, multiple breeding or agronomic experiments must be established within a limited field area during the same growing season. Traditionally, determining whether sufficient space remains for an additional trial may require manual inspection of field maps, spreadsheets, or communication among different research groups. By integrating trial allocation directly into Breedbase, the tool provides a centralized visualization of the trials that have already been placed within a location and the remaining available experimental area.

The visual representation of field occupancy enables researchers and research station managers to rapidly assess space availability before planning additional trials. Users can identify how many experiments have already been allocated, evaluate their spatial distribution, and determine whether sufficient contiguous space is available for a new experimental design. Consequently, the tool can improve coordination among research teams, reduce potential conflicts in field allocation, and support more efficient use of research station resources.

Furthermore, the integration of the Multi-Trial Allocation Tool with Breedbase connects spatial field planning with existing breeding and experimental information. Locations, seasons, and trials can be managed within the same breeding data management ecosystem, reducing the need for external planning tools and minimizing duplication of information. The interactive and flexible nature of the tool provides a practical approach for planning complex field research environments in which multiple experiments, researchers, and breeding programs share limited experimental space.

Beyond trial allocation, the tool also supports the generation of location-wide planting instructions for multiple trials (Figure 14). Planting parameters can be defined for the research location, including the number of rows per plot, number of plants per plot, plant spacing, row spacing, alleys between plots, and planting direction. Optional planting notes can also be included to provide additional instructions to field personnel. Importantly, these parameters are used to generate a consolidated planting map that considers the spatial arrangement of all trials allocated within the location. Consequently, field operations can be planned and executed using a unified representation of the entire research station rather than separate planting maps for each individual experiment.

**Figure 14:**
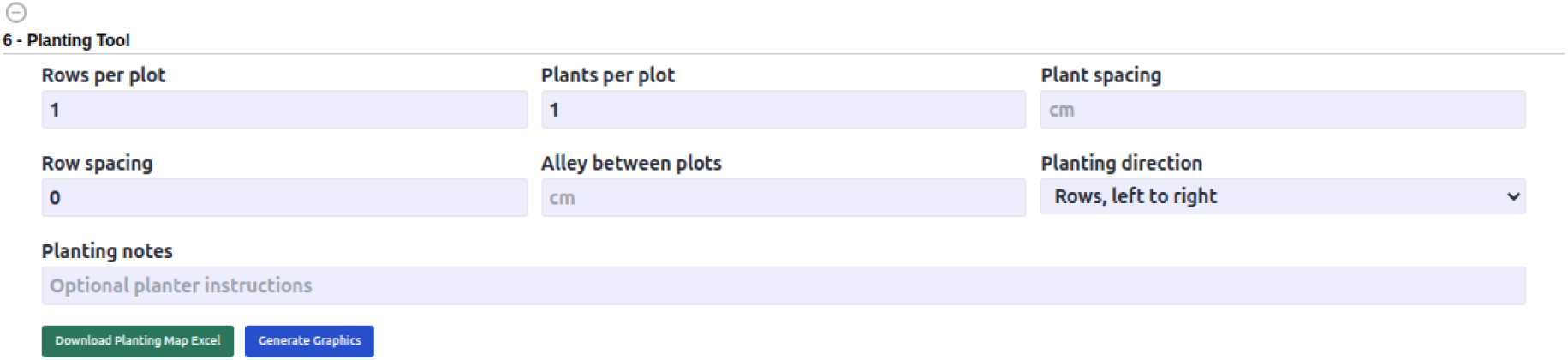
Planting tool settings for generating a planting map and defining field layout occupancy.

The planting direction and spatial position of the trials are incorporated into the generated field output, allowing the planting sequence to follow the actual organization of the location. The tool can produce an Excel-based planting map for use by field teams and generate graphical representations of the planned field layout. This functionality provides a connection between experimental design, spatial allocation, and field implementation, reducing the need to manually combine planting information from multiple trials. For research stations managing numerous experiments, a consolidated planting file may simplify field logistics, provide clearer instructions to planting teams, and reduce errors associated with plot identification and planting order.

By integrating multi-trial spatial planning and location-wide planting outputs directly into Breedbase, the tool links field logistics with existing breeding and experimental data management workflows. Locations, seasons, trials, field layouts, and planting instructions can be managed within the same information system, minimizing the duplication of data across independent planning tools. The combined functionality provides a practical framework for research stations in which multiple experiments, researchers, and breeding programs must share and coordinate the use of limited experimental space.

## Personalized Report Generator

The Personalized Report Generator was developed to address reporting and data validation requirements that are specific to individual Breedbase instances and breeding programs. Although Breedbase provides a broad set of standardized data access, visualization, and export tools, research institutions frequently require reports that reflect their own operational workflows, naming conventions, quality-control procedures, and data management practices. These requirements can differ substantially among crops, institutions, and breeding programs and may therefore be difficult to accommodate through a single predefined reporting interface. The personalized reporting framework provides a flexible mechanism for addressing these instance-specific needs without requiring every specialized report to become part of the general Breedbase reporting workflow.

Breedbase instances can request a customized report by defining the information, filtering criteria, validation rules, or output required by their research program. These requirements can be communicated to the Breedbase development team, which can implement a report-specific script according to the requested specifications. Personalized reports are implemented through Perl scripts that interact with the Breedbase data infrastructure and apply customized data retrieval and processing logic. This approach allows a report to combine information from different components of the database and to incorporate institution- or program-specific rules that may not be applicable to other Breedbase instances.

An example of a personalized report is the automated verification of whether accession names follow the naming convention established by a breeding program. Breeding programs commonly adopt specific nomenclature rules to encode information such as program identifiers, crossing years, population codes, selection cycles, or germplasm categories within accession names. As databases increase in size and accessions are created by multiple users or imported from different data sources, deviations from these conventions may occur and can reduce data consistency. A customized report can retrieve accessions associated with a breeding program, evaluate each accession name according to the program-specific naming rules, and identify records that do not conform to the expected format. The resulting report allows breeders and database curators to rapidly review potential naming inconsistencies and perform corrective actions when necessary. Because naming conventions may differ considerably among institutions and breeding programs, this type of validation is particularly well suited to the personalized reporting framework.

The framework can also support other complex reporting and data quality-control workflows. For example, customized reports may be developed to inspect phenotype-associated properties, identify inconsistencies in seedlot or stock names, summarize trial performance, or combine trial information, phenotypic observations, germplasm records, and experimental metadata according to institution-specific criteria. The report logic can therefore be tailored not only to retrieve information but also to evaluate, validate, summarize, and organize data according to the operational requirements of a particular Breedbase instance.

Once a personalized report script has been implemented and configured for an instance, it becomes available through the Personalized Report Generator interface (Figure 15). Authorized users can select one or more of the report scripts available for that database and, when applicable, provide parameters such as a start and end date. The reporting interface therefore provides a common entry point for executing multiple customized reporting workflows while allowing each report to retain its own specialized retrieval, validation, and processing logic. New report scripts can be added as additional requirements emerge, enabling the reporting capabilities of an individual Breedbase instance to evolve together with the needs of its users.

**Figure 15:**
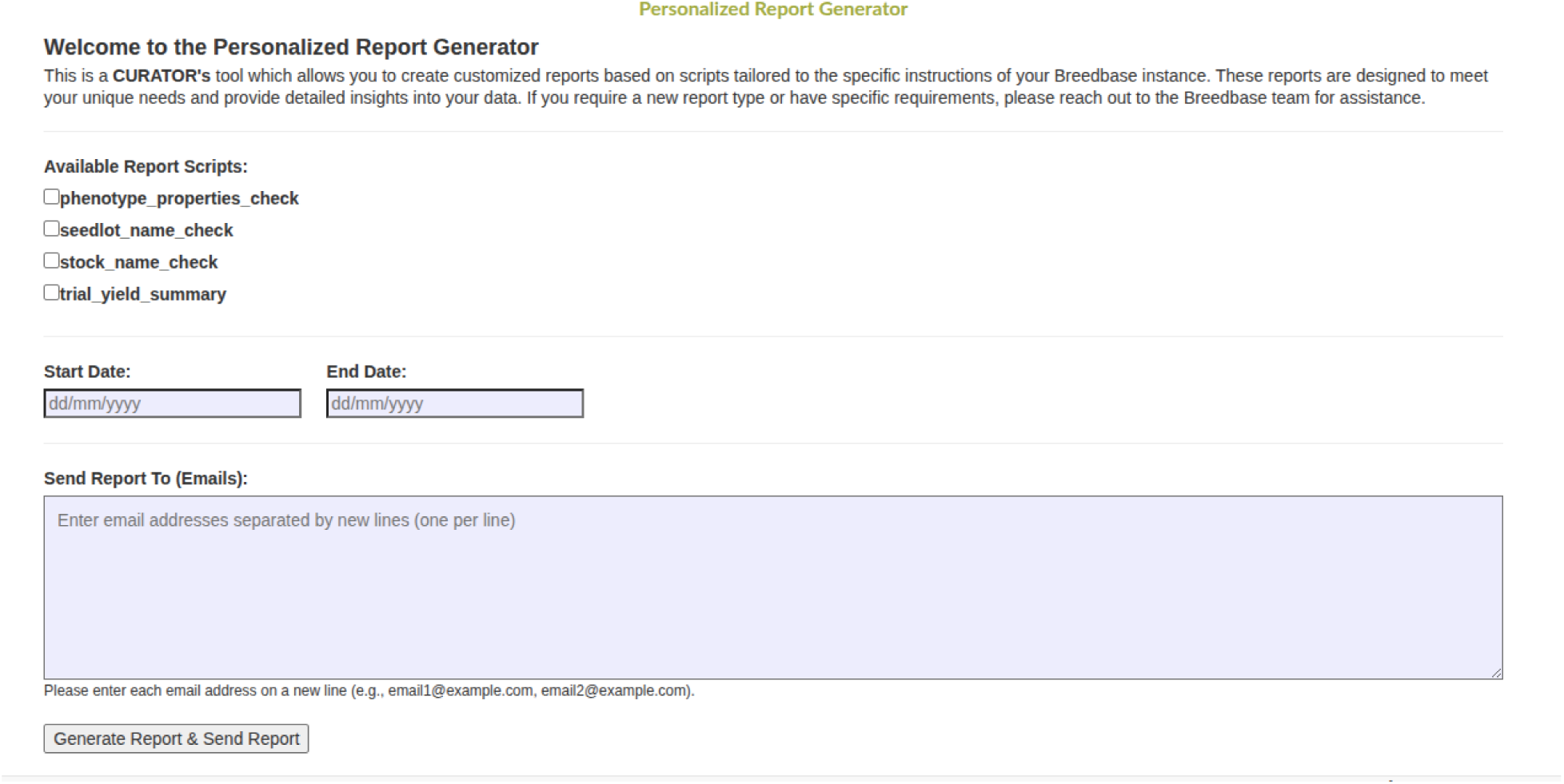
Personalized report generator showing scripts defined in the configuration file and email fields for analysis recipients.

Generated reports can also be distributed directly to one or more specified email recipients. Users can provide the email addresses of researchers, curators, or program managers who should receive the report, and the selected reporting workflows can generate and deliver the resulting information to the intended recipients. Direct report distribution can simplify collaborative data review and quality-control processes, particularly when the same information must be examined by multiple members of a research or breeding team. This functionality reduces the number of manual steps required to retrieve the data, prepare the output, and distribute the report among collaborators.

An important characteristic of the Personalized Report Generator is the separation between general Breedbase functionality and instance-specific reporting logic. Highly specialized reports may only be relevant to a single institution, crop, or breeding program and could introduce unnecessary complexity if incorporated into the standard reporting tools provided to all Breedbase users. By implementing such requirements as independent report scripts, Breedbase can preserve a broadly applicable core platform while still supporting highly customized institutional workflows. The reporting interface remains standardized, whereas the underlying data retrieval and validation logic can be adapted to the specific requirements of each Breedbase instance.

## Decision Meeting

The Decision Meeting tool was developed to support structured and data-informed decisions regarding the progression of breeding materials through a breeding program. A central component of the tool is the ability to assign a breeding stage to each accession independently within each breeding program. This program-specific stage assignment provides substantial flexibility for institutions in which the same accession may be evaluated, maintained, or utilized differently by multiple breeding programs. For example, an accession considered an advanced breeding line in one program may still be under preliminary evaluation in another program or may be retained as parental material for a different breeding objective. By associating breeding stages with the combination of accession and breeding program, rather than defining a single global stage for an accession, Breedbase can better represent the program-specific progression and use of germplasm.

This flexibility is particularly important in collaborative breeding environments, where germplasm is frequently shared among researchers, locations, and breeding programs. The value and developmental status of an accession are often dependent on the objectives of an individual program, target environment, market class, or trait of interest. Consequently, assigning a single breeding stage to an accession across an entire database may not accurately reflect its status in different research contexts. The decision framework implemented in Breedbase allows breeders to independently manage accession progression according to the selection strategy and advancement pipeline of each breeding program, while retaining the accession as a shared germplasm entity within the database.

The Decision Meeting tool provides a centralized interface for reviewing accessions and recording selection decisions (Figure 16). After creating a decision meeting, users can associate an existing accession list with the meeting and systematically evaluate the materials included in that list. For each accession, users can review its current breeding stage and record one of four primary decisions: Drop, Advance, Hold, or Jump. The *Drop* decision indicates that the accession should no longer progress in the current breeding pipeline, whereas *Advance* moves the accession to the next defined stage. *Hold* retains the accession at its current stage, allowing further evaluation before a progression decision is made. The *Jump* option provides additional flexibility by allowing an accession to bypass one or more intermediate stages and move directly to a later stage in the breeding process. This may be useful when exceptionally promising material is identified or when information from previous evaluations supports accelerated advancement.

**Figure 16:**
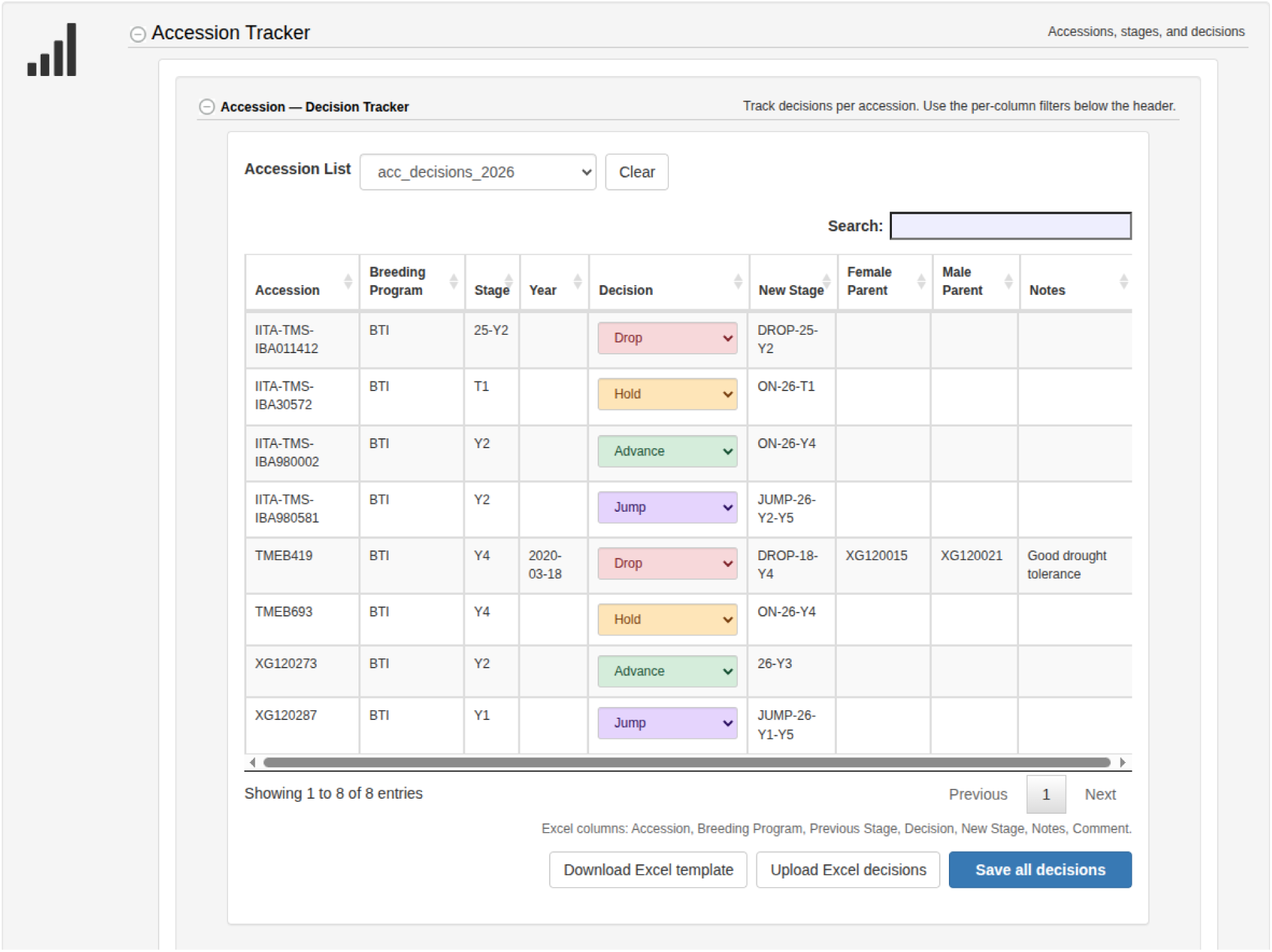
Decision meeting tracker, showing a list of accessions and decisions.

The use of explicit decision categories provides a standardized framework for recording outcomes from breeding review meetings. Rather than maintaining selection decisions in personal spreadsheets, written notes, or disconnected files, breeders can capture decisions directly within Breedbase and associate them with the corresponding accessions and breeding programs. The interface displays the previous stage, the recorded decision, and the resulting new stage, providing a transparent representation of how breeding material progresses through the pipeline. Importantly, both the decision categories and breeding stage names can be customized according to the configuration and workflow of each Breedbase database, allowing the tool to accommodate different breeding schemes, terminology, and advancement strategies across crops and breeding programs. Thus, the decision framework is not restricted to a predefined breeding pipeline but can be adapted to the operational structure of the institution using the database. Notes and additional contextual information can also be recorded to document the rationale for individual decisions, improving the traceability of the selection process and facilitating subsequent review by breeders and collaborators.

An additional objective of the Decision Meeting tool is to provide a historical and auditable record of the breeding selection process. Decisions made during each meeting are recorded, and the researchers or other participants involved in the decision process can be associated with the meeting. This information makes it possible to track not only how an accession progressed through the breeding pipeline, but also the context in which a decision was made and the individuals who participated in its evaluation. Such provenance is particularly valuable for long-term breeding programs, in which advancement or elimination decisions may need to be reviewed several years after the original selection meeting. By preserving decision histories and meeting participation, the tool supports greater transparency, institutional continuity, and traceability of breeding program management.

Breeding stage assignments are persistently stored in Breedbase and can be retrieved as accession properties. Consequently, the stage of an accession within a breeding program remains available beyond the individual decision meeting and can be accessed through other Breedbase data retrieval and management workflows. The preservation of stage information as an accession-associated property enables longitudinal tracking of germplasm progression and facilitates its use in downstream reporting, filtering, data extraction, and breeding program management.

To facilitate data-informed decision making, a phenotype dataset can also be associated with the decision meeting (Figure 17). Phenotypic information from the selected dataset is summarized and visualized within the decision environment, allowing users to evaluate accession performance while recording breeding decisions. Trait-specific graphical summaries, including boxplots and bar plots, provide an overview of the distribution and relative performance of accessions for selected traits. Users can select traits of interest and highlight one or more accessions in the visualizations, facilitating comparisons among breeding materials and supporting discussion during selection meetings. For example, the distribution of phenotypic observations for a target trait can be compared across accessions, providing information on both their relative performance and the variability observed in the available experimental data.

**Figure 17:**
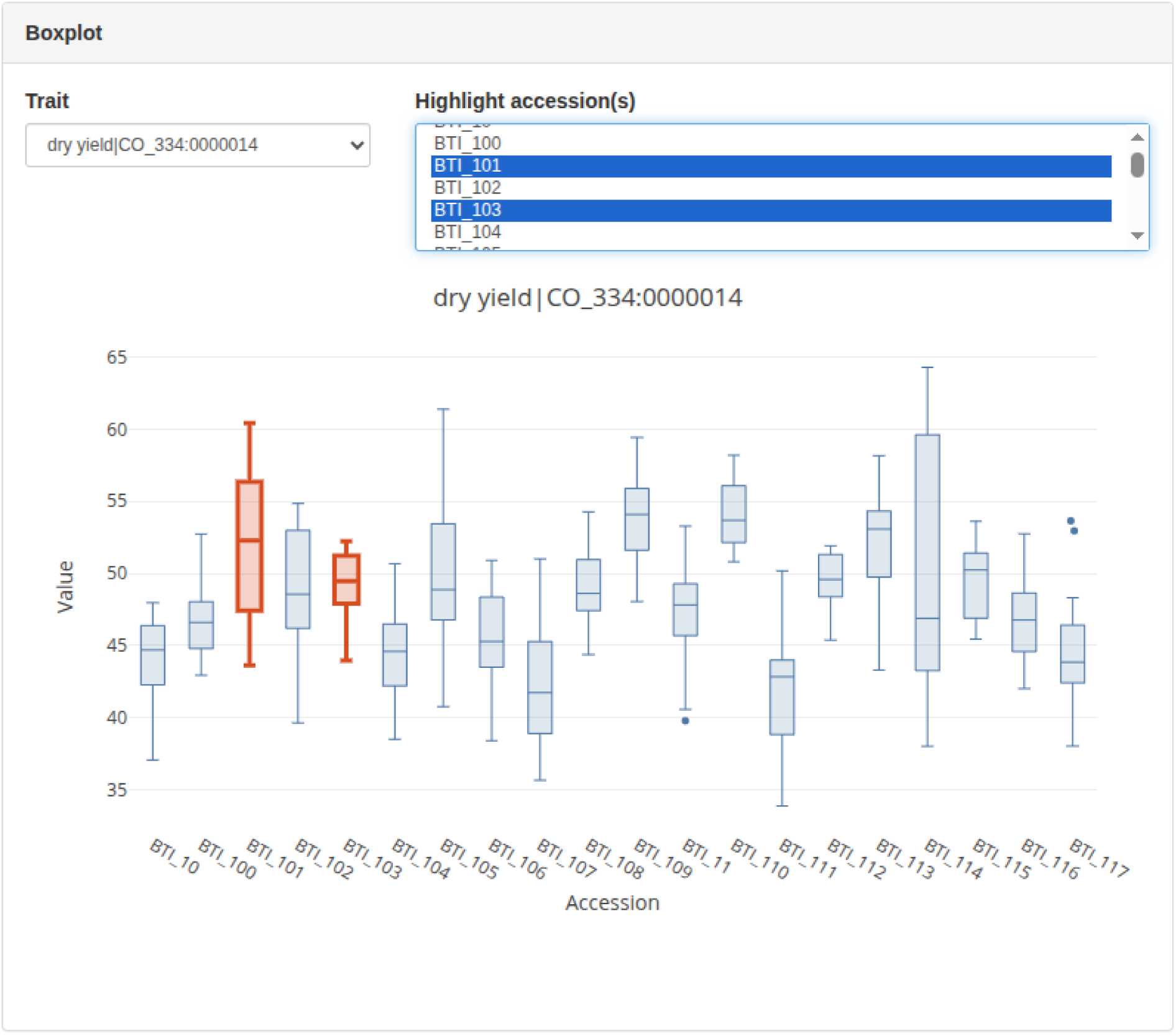
Boxplot of dry yield for selected accessions to support advancement decisions, highlighting accessions BT_101 and BTI_103.

The integration of phenotypic summaries with the decision interface creates a direct connection between experimental data and breeding pipeline management. Breeders can review performance information and record advancement decisions within the same workflow, reducing the need to alternate between external statistical outputs, spreadsheets, and breeding management systems. Although graphical summaries do not replace detailed statistical analyses or multi-environment evaluation, they provide an accessible overview of the available phenotypic data and can support rapid identification and discussion of promising or underperforming materials during breeding decision meetings.

For breeding programs that already conduct selection meetings using spreadsheets, the tool also supports the import of decisions from an Excel file. Users can download a structured Excel template, record or compile accession decisions externally, and subsequently upload the completed file into Breedbase. This functionality provides flexibility for different organizational workflows and facilitates the transition from spreadsheet-based decision management to an integrated breeding database. Decisions can therefore be entered interactively through the web interface or imported in batch, allowing breeding programs to select the approach that best fits their operational needs.

## Single Sign-On Logins

Authentication and authorization are critical components of any data sharing platform, in which the system verifies the user’s identity (authentication) and determines the resources the user may access (authorization). In Breedbase 1.0, users would create accounts by entering their preferred username and password, which was then stored and encrypted in the database using a cryptographic hash. Users could then access the system using a login form, which compared their supplied credentials with those stored in the database, and on a successful match, logged them into the database. This relatively simplistic model ensures full autonomy over the login process without depending on an external service, which provides clarity to users and simplicity in developing and troubleshooting.

However, internally managing user credentials comes with several disadvantages. A typical user has many web accounts that they must remember and manage passwords for, which becomes tedious and time-consuming. A consequence of this is that individuals may re-use passwords between accounts, which increases their risk should a data breach occur in any of the sites where the same password is used. This authentication approach also creates several obstacles that can impede a user accessing the system for the first time, as they must first successfully complete several login forms and email verification steps before accessing the database.

To address these problems, Breedbase 2.0 introduces a new Single Sign-On (SSO) option for users to login with an external identity provider. This has been implemented using the OAuth 2.0 protocol, specifically the Authorization Code Flow with OpenID Connect (OIDC). This standardized and highly secure protocol allows users to login with an external identity provider who may be a proprietary third party (ex. Google, Microsoft, ORCID) or a self-hosted service (ex. Keycloak, Authentik). When using the SSO feature, users will first be redirected to login through the provider’s page (ex. Google) and if that login is successful, will then be redirected back to the Breedbase application. These redirects occur in the user’s browser, and their SSO password is never sent to the Breedbase server.

Breedbase administrators can configure the connection to their preferred providers in a config file, and also determine if they trust the provider to ‘auto-provision’ new accounts. When this setting is enabled, new users may skip the account creation step, and instead have a new account automatically created for them after their identity is verified by the provider (ex. Google). This feature allows users to sign up and login with a single click, improving both ease of access and security in credential management. Administrators can also leverage additional security checks offered by the provider, such as Multi-Factor Authentication (MFA) to enforce stringent security measures for databases with highly sensitive information.

## Improved User Permission System

Breeding involves many different users, teams, and programs, all of whom manage data through the database instance, but may not require access to all the data in the database. In fact, it may be preferable that users managing genotyping data may not be able to change or even view phenotypic data, and vice versa. Particularly sensitive data, such as pedigree data, should only be viewable by a restricted set of users. In Breedbase 1.0, the permission system was relatively simple. There were only three levels of roles: The “user” role, which could read only; the “submitter” role, which could read and write to the database and modify their own submissions; and the curator role, which could read, write and modify anything in the database, including the user roles themselves.

In Breedbase 2.0, we have vastly expanded the capabilities of the permission system. The database has been sub-divided into about 20 different resource domains for each of which specific access parameters can be defined. The currently defined resources are phenotyping, genotyping, images, pedigrees, privileges, trials, crosses, wizard, community, loci, breeding_programs, stocks, catalog, user_roles, ontologies, publications, locations, seedlots, and the wiki system.

For each of these domains, any of the currently four defined access_levels can be defined for a role: read, write, update, and delete. The permissions are set as a combination of a role, a resource, and an access_level.

Permissions can be restricted to certain breeding programs or to the users themselves, by adding a requirement for breeding program membership or ownership of the data. By default, Breedbase 2.0 emulates the same permissions as Breedbase 1.0. However, roles can now be modified, and new roles can be added and freely defined. For example, a role “phenotyper” could be added with read and write access only to phenotypic information. Similarly, the user role “user” could be stripped of the ability to view pedigrees.

To set the permissions, a new page has been introduced, which allows to create the role / resource / access_level combinations. The page is available at /access/table, where currently defined permissions are displayed in a table format. Buttons for adding new permissions and new roles are available, and currently associated resources can be removed by clicking on an next to the resource. The breeding program and ownership restrictions are shown next to the resource name, as [BP] or [OWN]. The roles defined by this system can then be attributed to users with the same interface as in Breedbase 1.0.

## Wiki System

To simplify project management, a wiki system has been added so that many project related documents can be stored right in Breedbase, and do not require expensive subscriptions to other project management software that sometimes provide such or similar features. The wiki system is distinct from the Breedbase user manual (https://solgenomics.github.io/sgn/), which exists as a resource for users to understand the tools and workflows available on the website. Instead, the wiki system enables users to create their own documentation related to procedures, policies, and best practices they use in their breeding programs. A breeding program administrator could, for example, create wiki documentation detailing how to manage trait terms in their database or what data should be included in printed barcode labels created using the Label Designer. Breeding programs and laboratories can use this feature to organize and integrate their field protocols with Breedbase, serving the broader effort of making Breedbase a cornerstone component of the digital agriculture ecosystem. The wiki system is available at /wiki and is available to logged in users. Access restrictions can be defined by the new permission system, but by default, the “user” role can only view, the “submitter” role can view and create pages, as well as modify the pages created, whereas the curator view can view, create, and modify any page.

## Transformation and Tracking Re-generation

While Breedbase was initially designed for traditional breeding and genomic prediction, transgenics remain an important technology for both agriculture and plant science (Anwar and Kim 2020). A system has been implemented that can manage the generation of transgenic plants, and track the process from transformation to regenerated plants, which can then be evaluated in the same way as traditional accessions in the Breedbase system, such as in field trials.

Transformation events can be recorded in the database by creating a transformation project, which will group related transformation events. One or more accessions to be transformed need to be available in the database, as well as one or more vectors. The vector type was introduced as an additional stock type, with vectors having additional attributes, describing such features as the vector backbone, inserted genes, and selection markers. Vectors can also be viewed graphically if a well-formed Genbank formatted file describing the vector is uploaded.

A transformation is modelled similarly to a cross, with the untransformed accession being the female_parent and the vector standing in for the male_parent accession, but with a relationship type of transformant_of. In the pedigree view, the transformation event appears as an additional relationship.

From a successful transformation event, many new accessions can be generated; new transformant accessions can be added for each experiment with a simple interface. Naming schemes can be defined in the database which will be used to name new transformants automatically. New transformants are usually screened for presence of insertions, insertion number, and expression levels. These results can be stored in the database and can facilitate the selection of appropriate transgenic germplasm for downstream use.

During the regeneration process, progress can be tracked using a tablet using a specific webpage in the database that can read barcodes through the tablet’s camera. When a barcode is read, the page for that specific sample will open, displaying all the previous collected data, if available. Tracking data can be entered for each transformation event using a custom vocabulary, which can be independently defined for each instance of Breedbase. For example, the vocabulary could have entries for medium changes, rooting events, and plantlet counts.

The regeneration tracking is also available independently of the transformation tool and can be used to track such processes as embryo rescue or tissue regeneration experiments. By expanding functions to plant transformation and transgenics, Breedbase should appeal more to research labs and advanced breeding programs. More details on the transformation system will be published in a separate paper.

## Discussion

Breedbase is an open source breeding database offering a wide and growing variety of features and tools, making it a feature-rich platform that can be adapted to most breeding and plant research needs. Indeed, Breedbase instances exist for crops as varied as cassava, sweet potato, yam, banana, blueberries, potato, tomato, maize, wheat, barley, and many more.

Over the last few years, as described in this paper, we have added many features, including the improved handling of repetitive measurements; a redesigned experimental treatments interface; a more fine-grained user permission system; improved trial quality control workflows, intercropping support; the addition of a “decision meeting” feature that helps decision makers make the best selection with the available data; a field-level trail layout tool; a single sign-on login to leverage authentication providers such as Google and ORCID; tracking of plant transformation and transgenics; and an integrated wiki system.

Most of these features were added because users requested them, and user feedback and guidance has been taken into account through all stages of development. Users can submit tickets to the GitHub account to request new features or bug fixes. Currently, the issue tracker contains over 500 open tickets, mostly of the feature_request type, with more than 3000 having been closed. Based on these tickets, new features will be prioritized and will continue to be added in the future, making the Breedbase system easier to use and more useful to more breeding programs over time.

Artificial intelligence (AI) brings new opportunities for both research and breeding, as AI has the promise to speed up software development by a huge factor.(Vaswani et al. 2017) AI-enabled databases will also be much easier to use, as AI can provide a natural language interface to query the database. It also has the potential to vastly improve the predictions of traits, genome annotation, and other novel tools for making breeding more efficient. We are currently working on this next set of technologies, which will be the subject of another paper. However, the importance of Breedbase and similar databases will always be the collection and safeguarding of ground truth data for crop improvement and research, and to democratize the complex data management that breeding requires through open source tools.

## Author Contributions

Roles follow the CRediT taxonomy (name first, roles following), in manuscript author order.

**Srikanth Kumar Karaikal** - Writing—original draft, Writing—review & editing

**Ryan Preble -** Formal analysis, Visualization, writing—original draft, Writing—review & editing

**David J Waring -** Writing—original draft, Writing—review & editing

**Christiano C Simoes -** Conceptualization, Writing—original draft, Writing—review & editing

**Naama Menda -** Investigation, writing—review & editing

**Titima Tantikanjana -** Writing—review & editing

**Jing Zhang –** Investigation

**Katherine Eaton -** Writing—review & editing

**Keo Corak –** Investigation

**Clayton Birkett -** Investigation, conceptualization

**Allison Krill Brown -** Writing—review & editing

**Mitchell Feldmann -** Investigation, conceptualization

**Christian Lamm -** Writing—review & editing

**Mercedes Thieme -** Writing—review & editing

**Prasad Peteti -** Conceptualization and investigation

**Heather Manching -** Visualization and validation

**Adrian Powell -** Investigation,

**Jean-Luc-Jannink -**Funding acquisition

**Lukas A Mueller -** Writing—original draft, Writing—review & editing, Funding acquisition

## Acknowledgments

We would like to thank the breeding programs, their leaders, their staff, and funders, for providing so much feedback and so many suggestions, particularly the CGIAR centers IITA, CIAT, CIP, CIMMYT, and IRRI, as well as the many National Programs such as NRCRI, NaCRRI, TARI, and many others. We would also like to thank users who took the effort to write GitHub tickets for the issues they encountered, and those who helped with quality control of the code.

## Conflicts of Interest

No conflicts of interest

## Funding

Breedbase has been funded over the years by many different funding agencies: The National Science Foundation (NSF), the United States Department of Agriculture (USDA), the Bill and Melinda Gates Foundation (BMGF), and Gates AgOne. We thank the funding agencies for their support.

## Data availability

All breedbase code and resources are freely available for download and reuse under the MIT license. Software is hosted at https://github.com/solgenomics/sgn and up-to-date site images are available at https://hub.docker.com/u/breedbase/.

## Notes

### Competing Interest Statement

The authors have declared no competing interest.

## References

Andrews DJ, Kassam AH. 1976. The Importance of Multiple Cropping in Increasing World Food Supplies. In: Multiple Cropping. John Wiley & Sons, Ltd; p 1–10 [accessed 2026 May 12]. https://onlinelibrary.wiley.com/doi/abs/10.2134/asaspecpub27.c1. 10.2134/asaspecpub27.c1

Anwar A, Kim J-K. 2020. Transgenic Breeding Approaches for Improving Abiotic Stress Tolerance: Recent Progress and Future Perspectives. Int J Mol Sci. 21(8):2695. 10.3390/ijms21082695

Aparicio J et al. 2024. Mr.Bean: a comprehensive statistical and visualization application for modeling agricultural field trials data. Front Plant Sci. 14 [accessed 2026 May 5]. https://www.frontiersin.org/journals/plant-science/articles/10.3389/fpls.2023.1290078/full. 10.3389/fpls.2023.1290078

Ashburner M et al. 2000. Gene Ontology: tool for the unification of biology. Nat Genet. 25(1):25–29. 10.1038/75556

Breseghello F, Coelho ASG. 2013. Traditional and modern plant breeding methods with examples in rice (Oryza sativa L.). J Agric Food Chem. 61(35):8277–8286. 10.1021/jf305531j

Brooker RW et al. 2015. Improving intercropping: a synthesis of research in agronomy, plant physiology and ecology. New Phytologist. 206(1):107–117. 10.1111/nph.13132

Cobb JN et al. 2019. Enhancing the rate of genetic gain in public-sector plant breeding programs: lessons from the breeder’s equation. Theor Appl Genet. 132(3):627–645. 10.1007/s00122-019-03317-0

Conway G. 1998. The Doubly Green Revolution: Food for All in the Twenty-first Century. Cornell University Press.

Crossa J et al. 2017. Genomic Selection in Plant Breeding: Methods, Models, and Perspectives. Trends Plant Sci. 22(11):961–975. 10.1016/j.tplants.2017.08.011

Dumschott K et al. 2023. Ontologies for increasing the FAIRness of plant research data. Front Plant Sci. 14:1279694. 10.3389/fpls.2023.1279694

Evenson RE, Gollin D. 2003. Assessing the impact of the green revolution, 1960 to 2000. Science. 300(5620):758–762. 10.1126/science.1078710

Feldmann MJ, Pincot DDA, Cole GS, Knapp SJ. 2024. Genetic gains underpinning a little-known strawberry Green Revolution. Nat Commun. 15(1):2468. 10.1038/s41467-024-46421-6

Hale I et al. 2025. Sustaining public plant breeding programs across generations. Crop Science. 65(3):e70094. 10.1002/csc2.70094

Hershberger J et al. 2021. Making waves in Breedbase: An integrated spectral data storage and analysis pipeline for plant breeding programs. The Plant Phenome Journal. 4(1):e20012. 10.1002/ppj2.20012

Holmes GJ. 2024. The California Strawberry Industry: Current Trends and Future Prospects. International Journal of Fruit Science. 24(1):115–129. 10.1080/15538362.2024.2342900

Jannink J-L, Lorenz AJ, Iwata H. 2010. Genomic selection in plant breeding: from theory to practice. Brief Funct Genomics. 9(2):166–177. 10.1093/bfgp/elq001

López-Bellido L, Fuentes M, Castillo JE, López-Garrido FJ. 1998. Effects of tillage, crop rotation and nitrogen fertilization on wheat-grain quality grown under rainfed Mediterranean conditions. Field Crops Research. 57(3):265–276. 10.1016/S0378-4290(97)00137-8

López-Bellido L, López-Bellido RJ, Redondo R. 2005. Nitrogen efficiency in wheat under rainfed Mediterranean conditions as affected by split nitrogen application. Field Crops Research. 94(1):86–97. 10.1016/j.fcr.2004.11.004

Menda N et al. 2025. Post-composing ontology terms for efficient phenotyping in plant breeding. Database (Oxford). 2025:baaf020. 10.1093/database/baaf020

Meuwissen TH, Goddard ME. 2001. Prediction of identity by descent probabilities from marker-haplotypes. Genet Sel Evol. 33(6):605–634. 10.1186/1297-9686-33-6-605

Moore VM, Peters T, Schlautman B, Brummer EC. 2023. Toward plant breeding for multicrop systems. Proc Natl Acad Sci USA. 120(14):e2205792119. 10.1073/pnas.2205792119

Morales N et al. 2022. Breedbase: a digital ecosystem for modern plant breeding. G3 Genes|Genomes|Genetics. 12(7):jkac078. 10.1093/g3journal/jkac078

Plucknett DL, Smith NJH. 1986. Sustaining Agricultural Yields. BioScience. 36(1):40–45. 10.2307/1309796

Ribaut J-M, Hoisington D. 1998. Marker-assisted selection: new tools and strategies. Trends in Plant Science. 3(6):236–239. 10.1016/S1360-1385(98)01240-0

Rife TW, Poland JA. 2014. Field Book: An Open-Source Application for Field Data Collection on Android. Crop Science. 54(4):1624–1627. 10.2135/cropsci2013.08.0579

Selby P et al. 2019. BrAPI—an application programming interface for plant breeding applications. Bioinformatics. 35(20):4147–4155. 10.1093/bioinformatics/btz190

Selby P et al. 2025. BrAPI v2: real-world applications for data integration and collaboration in the breeding and genetics community. Database (Oxford). 2025:baaf048. 10.1093/database/baaf048

Shrestha R et al. 2012. Bridging the phenotypic and genetic data useful for integrated breeding through a data annotation using the Crop Ontology developed by the crop communities of practice. Front Physiol. 3:326. 10.3389/fphys.2012.00326

Smith B et al. 2007. The OBO Foundry: coordinated evolution of ontologies to support biomedical data integration. Nat Biotechnol. 25(11):1251–1255. 10.1038/nbt1346

Tecle IY et al. n.d. Breedbase: An Integrated Breeding Informatics System with Statistical Analysis and Decision-Support Tools.

Thapa R et al. 2024. Remote sensing for estimating genetic parameters of biomass accumulation and modeling stability of growth curves in alfalfa. G3 Genes|Genomes|Genetics. 14(11):jkae200. 10.1093/g3journal/jkae200

Tilman D et al. 2002. Agricultural sustainability and intensive production practices. Nature. 418(6898):671–677. 10.1038/nature01014

Vandermeer JH. 1989. The Ecology of Intercropping. Cambridge University Press. [accessed 2026 May 12]. https://www.cambridge.org/core/books/ecology-of-intercropping/ACD6773C1D43F28D8B499F5D36A649D1. 10.1017/CBO9780511623523

Vaswani A et al. 2017. Attention is All you Need. In: Advances in Neural Information Processing Systems. Vol. 30. Curran Associates, Inc. [accessed 2026 June 30]. https://proceedings.neurips.cc/paper_files/paper/2017/hash/3f5ee243547dee91fbd053c1c4a845aa-Abstract.html

Wallace JG, Rodgers-Melnick E, Buckler ES. 2018. On the Road to Breeding 4.0: Unraveling the Good, the Bad, and the Boring of Crop Quantitative Genomics. Annu Rev Genet. 52:421–444. 10.1146/annurev-genet-120116-024846

